# Loss of genetic variation in the two-locus multiallelic haploid model

**DOI:** 10.1101/2020.03.07.981852

**Authors:** Martin Pontz, Marcus W. Feldman

## Abstract

In the evolutionary biology literature, it is generally assumed that in deterministic haploid selection models, in the absence of variation-generating mechanisms such as mutation, no polymorphic equilibrium can be stable. However, results corroborating this claim are scarce and almost always depend upon additional assumptions. Using ideas from game theory, we establish a condition on the fitness parameters of haplotypes formed by two loci such that a monomorphism is a global attractor. Further, we show that no isolated equilibrium exists, at which an unequal number of alleles from two loci is present. Under the assumption of convergence of trajectories to equilirium points, we settle the two-locus three-allele case for a fitness scheme formally equivalent to the classical symmetric viability model.

## Introduction

A recent paper by Novak and Barton (2017) raises one of the main questions of population genetics right in the title: “When does frequency-independent selection maintain genetic variation?” They note that, while the answer is generally assumed to be “never” for constant selection acting on an idealized haploid population, basically only the cases of no recombination and of no selection have been solved and corroborate this claim. Well known results from perturbation theory, of course, expand these results to small parameter values of the respective force. In fact, Novak and Barton (2017) give a new, more standard, proof for the case of weak selection. The other extreme, tight linkage (low recombination rate), was solved by Kirzhner and Lyubich (1997), who also arrive at the same conclusion for additive fitnesses and arbitrary linkage. These three results hold for any number of loci and any number of alleles. They all incorporate convergence of the solutions to equilibrium points, via the powerful method of identifying a Lyapunov function.

Besides the general additive case and the trivial one-locus case, only the two-locus two-allele case has recieved attention for intermediate values of recombination and selection. Feldman (1971) was one of the first to rigorously analyze existence and stability of polymorphic equilibria in a two-locus two-allele haploid system with a simple fitness scheme. He showed that whenever a polymorphism exists, it is unique and unstable. A general fitness scheme was considered by Rutschman (1994). He showed convergence of the trajectories to equilibrium points in most parameter regimes. However, parameter combinations in which an internal (polymorphic) equilibrium was possible, couldn’t be treated in the same way. The final answer to the question of loss of genetic variation in the two-locus two-allele case is the paper by Bank, Bürger and Hermisson (2012). They showed that for the fitness parameter combinations not covered by Rutschman, an equilibrium exists, but it is always unstable.

We consider a well mixed haploid population with constant selection on two loci, each with an arbitrary number of alleles. Our fitness scheme is general without any restriction on the epistatic interaction between alleles. For convenience, the dynamics are stated in continuous time. First, we apply a method used on a game theoretic problem by Hofbauer and Su (2016). With its help we can show that if one allele dominates all the others from the same locus, then this allele becomes fixed. An allele dominating another here means that the fitness of an haplotype containing the *dominating* allele is greater than the fitness of an haplotype containing the other, *dominated*, allele for every choice of background allele. Further, we state and prove that no isolated equilibrium exists if the numbers of alleles at the two loci are unequal. This is done by finding a system of linear equations, whose solution corresponds to an internal equilibrium. For unequal numbers of alleles at the two loci, this system is overdetermined and thus, by basic linear algebra has, in general, no solution.

We also use the above-mentioned system of linear equations to give a different proof of the result by Bank, Bürger and Hermisson (2012). Finally, we show that there is a unique unstable polymorphic equilibrium in the two-locus three-alleles model with centrosymmetric fitnesses.

## Model

In the two-locus haploid model considered here, we assume that at one locus the alleles are *A*_1_, …, *A*_*m*_, while at the other locus the alleles are *B*_1_, …, *B*_*n*_. Let *p*_*ij*_ and *s*_*ij*_ be the frequency and the fitness, respectively, of haplotype *A*_*i*_*B*_*j*_ and define the matrices *S* = (*s*_*ij*_)_*m*×*n*_ an *P* = (*p*_*ij*_)_*m*×*n*_. In the following, we will mainly use the vector *p*, which is defined as the vector of all rows of *P*, i.e., *p* = (*p*_11_, …, *p*_1*n*_, *p*_21_, …, *p*_*m*1_, …, *p*_*mn*_)^*T*^. Following Nagylaki (1992, pp. 189–195) and Novak and Barton (2017), we can write the change in frequency over time, 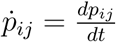, as

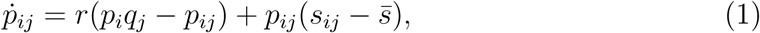

where 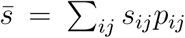 is the mean fitness, 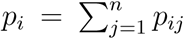 and 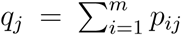 are the marginal frequencies of the alleles. As always, the sum of all haplotype frequencies is one, ∑_*ij*_ *p*_*ij*_ = 1. The quantities *p*_*i*_*p*_*j*_ − *p*_*ij*_ are measures of linkage disequilibrium (LD), and *r* > 0 denotes the recombination rate.

We investigate stability properties of the monomorphic equilibria and existence and stability properties of polymorphic equilibria. At equilibrium, the coordinates are denoted by a^ and satisfy

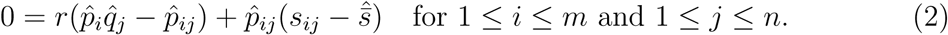

These are *mn* quadratic equations in *mn* variables.

The state space of (1) is the *mn*−dimensional simplex Δ_*mn*_ as defined by

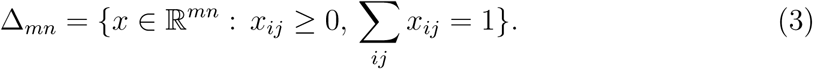

### Remark 1.

*We note that system* (1) *is invariant with respect to adding a constant c to S. If* 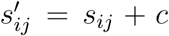, *then* 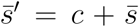 *and thus* 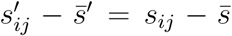 *for every pair* (*i, j*). *It will be especially useful to take* 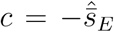, *where* 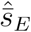 *denotes mean fitness at an equilibrium E. At the same equilibrium in the scaled fitness scheme the scaled mean fitness is* 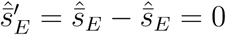.

## Results

### Stability of monomorphic equilibria

First, we determine the conditions under which monomorphism are stable, which is in turn used to give an upper bound for the number of stable hyperbolic monomorphic equilibria.

It is easy to see that each monomorphism is an equilibrium for (1).

To determine the local stability of equilibria, we compute the Jacobian *J* of 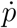, given by (1). For every pair (*i, j*) the following holds:

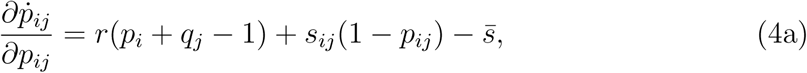

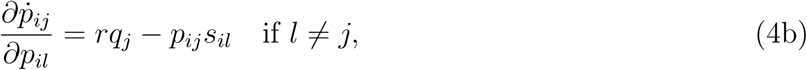

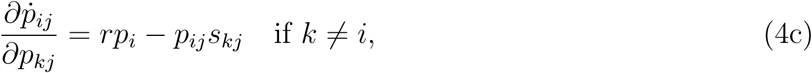

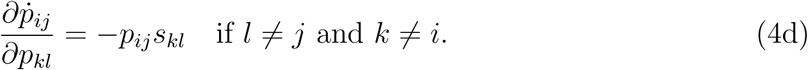

If we fix (*u, v*) and sum over the corresponding column of *J*, we get:

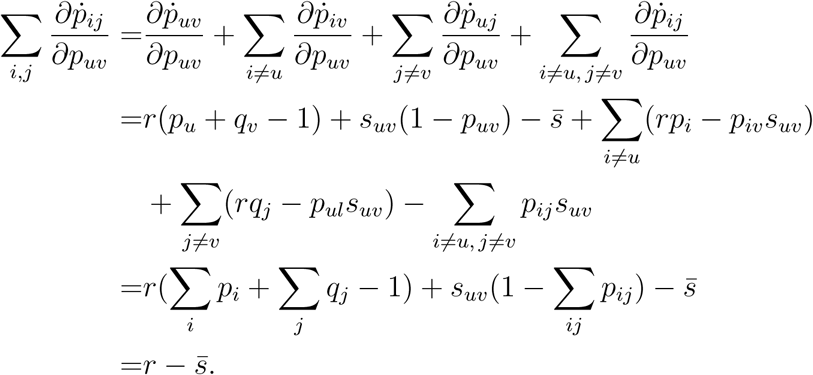

Since this holds for every column of the Jacobian, (1, …, 1) is a left eigenvector with the eigenvalue 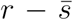. Because this eigenvector is normal to the simplex, the corresponding eigenvalue carries no information about the stability of an equilibrium.

Keeping this in mind, we can compute the Jacobian at the monomorphism with *p*_11_ = 1. At this equilibrium, 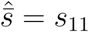. Applying this to (4a) yields 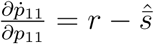.

The other diagonal entries are given by

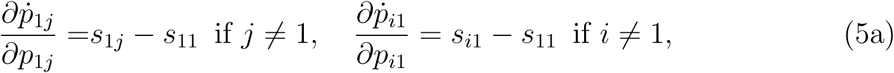

and

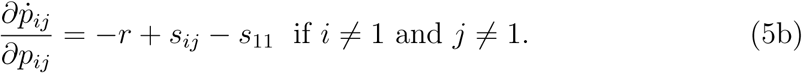

The remaining non-zero entries are given by:

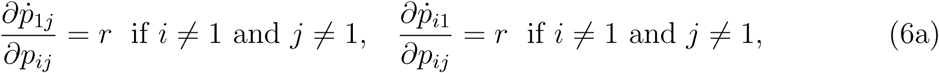

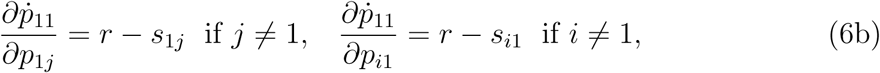

and

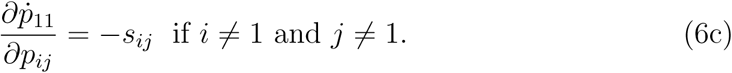

All other entries are zero.

After recalling the lexicographical order of the double indices in the vector *p*, inspection of the non-zero entries shows that 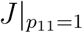 is an upper right triangular matrix and therefore, the eigenvalues are the diagonal entries given by (5).

We can, in general, relabel alleles and loci, such that the Jacobian of any monorphism is an upper right triangular matrix. This means that for the monomorphism 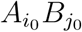, the *mn* − 1 eigenvalues that determine stability are given by

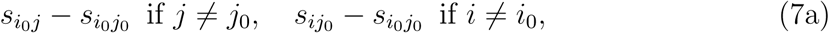

and

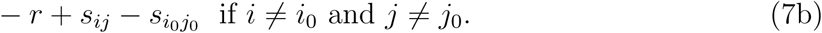

However, for certain choices of the fitness values, some of the monomorphisms are not isolated and have an eigenvalue equal to zero as the following result shows.

#### Lemma 1.

*Every point on the edge connecting* 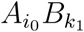 *with* 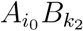, *given by* 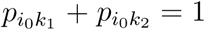, *is an equilibrium if and only if* 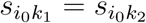.

*Proof.* On the edge connecting 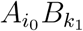 with 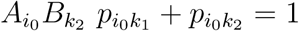 and *p*_*ij*_ = 0 for all *i* and *j* that do not form the pairs (*i*_0_*k*_1_) or (*i*_0_*k*_2_). Hence, *p*_*i*_*q*_*j*_ = 0 and thus, after plugging this into (1), 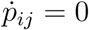 if *i* ≠ *i*_0_ and *j* ≠ *k*_1_, *k*_2_. Therefore, the only equations of (1) with non-zero right-hand side are

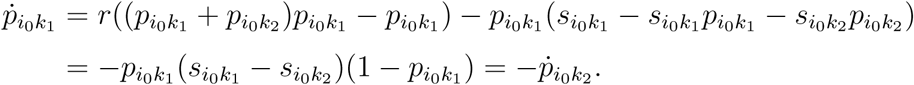

This is zero for all 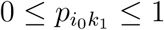 if and only if 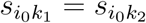. □

#### Remark 2.

*An analoguos result holds for the edge connecting* 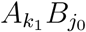 *with* 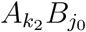

To avoid the complications of degenerate cases, we assume for this and the next section that

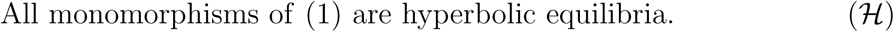

This means that no eigenvalue of a monomorphic equilibrium is zero. In particular, this implies that no two fitness values of haplotypes that share an allele are the same.

#### Proposition 1.

*The monomorphism* 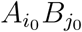 *is asymptotically stable if and only if* 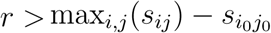. *In this case* (*i*_0_, *j*_0_) *is the unique pair such that* 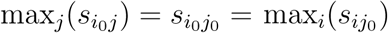.

*Proof.* The eigenvalues of the monomorphism 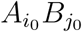 given by (7) are negative if and only if (*i*_0_*j*_0_) is the unique pair such that 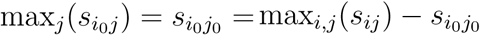 and 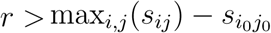 holds. □

The number of asymptotically stable monomorphisms is bounded by the following

#### Corollary 1.

*In a system with n alleles present at one locus and m at the other, there are between one and* min(*m, n*) *asymptotically stable monomorphisms.*

*Proof.* By Proposition 1, the monomorphism corresponding to the largest entry of the fitness matrix is always locally asymptotically stable because *r* > 0.

Let *m* < *n* and let *m* be the number of rows of the fitness matrix. Assumption (ℋ) implies that each row has a unique maximum. There are exactly *m* such maxima. If each of those is also the maximum in its respective column then the corresponding monomorphisms are locally asymptotically stable for sufficiently large *r* by Theorem 1. There cannot be more. □

#### Corollary 2.

*Let* 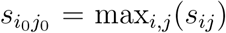, *then for every r* > 0 *the monomorphism* 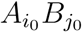 *is the only locally asymptotically stable monomorphism if and only if for all pairs* (*k, l*) *s*_*kl*_ ≤ max (max_*i*_(*s*_*il*_), max_*j*_(*s*_*kj*_)) *with equality only if* (*k, l*) = (*i*_0_, *j*_0_), *i.e.*, 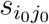 *is the only value that is maximal in both its row and column.*

*Proof.* For each pair (*k, l*) ≠ (*i*_0_, *j*_0_) assume that *s*_*kl*_ < max (max_*i*_(*s*_*il*_), max_*j*_(*s*_*kj*_)). Then by Proposition 1 and Corollary 1 it is clear that for every *r* > 0, 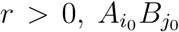 is the only locally asymptotically stable monomorphism.

Conversely, suppose there is a monomorphism *A*_*k*_*B*_*l*_ with *k* ≠ *i*_0_ and *l* ≠ *j*_0_ such that max_*j*_ *s*_*kj*_ = *s*_*kl*_ = max_*i*_ *s*_*il*_. There exists 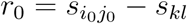. Then (7) implies that *A*_*k*_*B*_*l*_ is stable for *r* > *r*_0_. This contradicts the assumption that for every 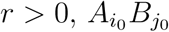 is the only stable monomorphism. □

If we assume that genetic variation is never maintained in a haploid population under selection and recombination, then the monomorphism described in Corollary 2 would be the natural candidate for a global attractor. However, no proof could be found to verify this.

### Dominating alleles

In the following, we apply ideas from game theory, in particular from a paper by Hofbauer and Su (2016) about dominating strategies, to alleles. For a special class of fitness schemes, this allows us to prove global stability of a monomorphism.

#### Theorem 1.

*If there exist alleles* 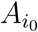 *and* 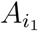 *such that* 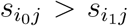 *holds for every j, then* 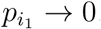.

*Proof.* Without loss of generality, suppose that 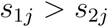 holds for all *j*. We show that the minimal quotient 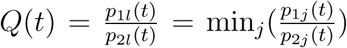 is increasing along trajectories in *t*. First, note that *l* = *l*(*t*) can assume different values in {1, …, *m*} at different times. Further, if *p*_1_, *p*_2_ > 0, then for every given *t*

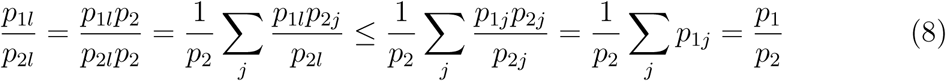

holds, which is equivalent to

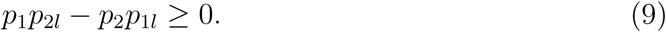

Then,

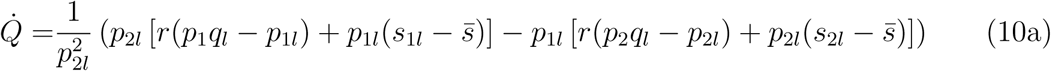

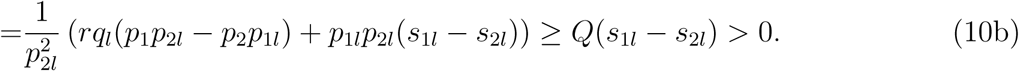

Define *δ* = min_*l*_(*s*_1*l*_ − *s*_2*l*_) > 0. Then (10) implies

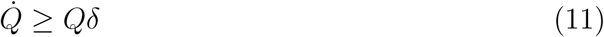

and subsequently

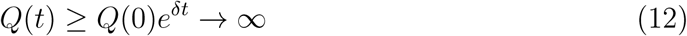

as *t* → ∞.

This implies that 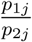 goes to infinity as *t* → ∞ for all *j*. Since the numerator is bounded, 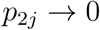 for all *j* and thus *p*_2_ → 0 as *t* → ∞. □

#### Remark 3.

*A similar theorem holds for the other locus.*

This can be interpreted such that allele 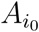 dominates 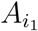, since for every background allele *B*_*J*_ the haplotype containing allele 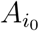 is fitter than that containing 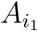. Almost intuitively, this leads to the loss of 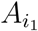. Ultimately, if 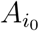 dominates all other alleles at the same locus, these should all go extinct. This is formalized in the following

#### Theorem 2.

*If there is an allele* 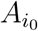 *with* 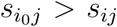 *for all i* ≠ *i*_0_ *and all j, then the monomorphism* 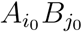 *with* 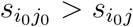 *for all j* ≠ *j*_0_ *is globally asymptotically stable.*

*Proof.* For each *i* ≠ *i*_0_ we apply Theorem 1. Then allele 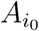 becomes fixed in the population. Because of (ℋ), there exists a unique *j*_0_ such that 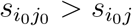 for all *j* ≠ *j*_0_. By (7) the monomorphism 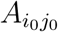 is the only asymptotically stable monomorphism □

### Polymorphic equilibria

Monomorphisms are equilibria where exactly one allele at each locus is present. Next, we look at equilibria where at least three alleles are involved.

While system (1) with state space Δ_*mn*_, with *m* < *n*, can always be imbedded in the system with state space Δ_*nn*_ where both loci have the same number of alleles, it simplifies derivations if we think of the equilibria in subsystems as being fully polymorphic.

From now on, we are thus mainly interested in the existence of fully polymorphic equilibria, i.e., states, where all haplotypes are present. Therefore, the inequalities

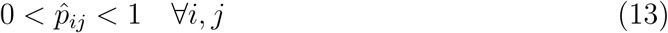

hold, which is equivalent to 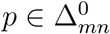. Here, ^0^ denotes the interior of the simplex.

Our main result is

#### Theorem 3.

*If m* ≠ *n, then* (2) *has either no or infinitely many solutions for which* (13) *holds. Thus, there are no isolated equilibria with all mn haplotypes present.*

In order to prove the theorem, we first need

#### Lemma 2.

*Define the matrix* 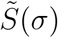 *by*

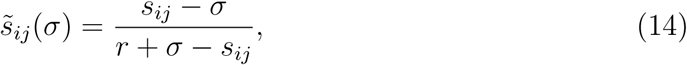

*where σ* ∈ ℝ. *For a given fitness matrix S and r* > 0, *a solution of* (2) *that fulfills* (13) *exists if and only if* 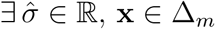 *and* **y** ∈ Δ_*n*_ *such that*

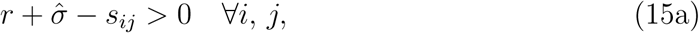

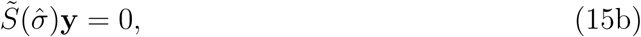

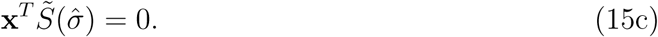

*If a solution exists, it is given by*

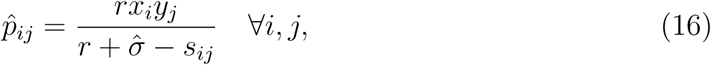

*and then*

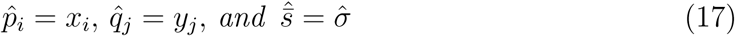

*holds.*

Here, Δ_*m*_ denotes the *m*−dimensional simplex.

*Proof.* (⇒) From (2), we can compute the following identities:

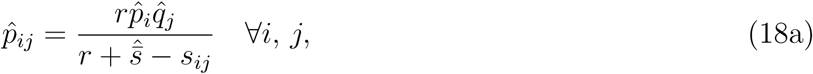

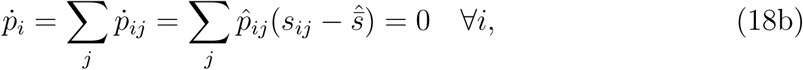

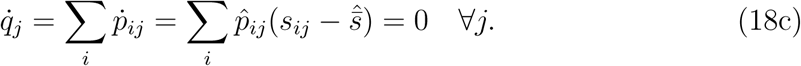

We plug (18a) into (18b) and (18c) to get:

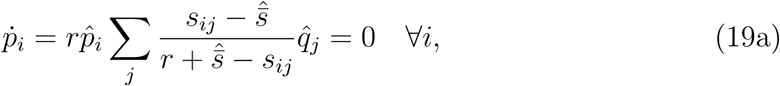

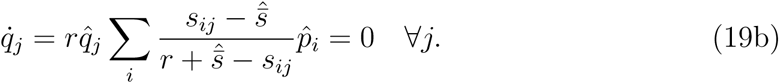

Then (13) entails 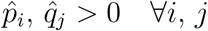. Thus, we can write (19) as

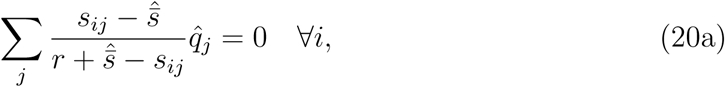

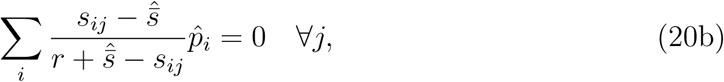

or in matrix terms, after we set 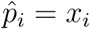 and 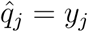:

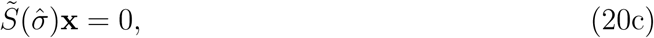

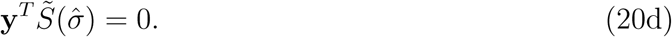

The remaining condition (15a) is implicit in (18a) because of (13).

(⇐) We define

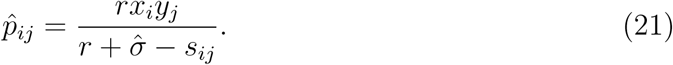

We have to show that

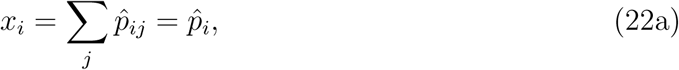

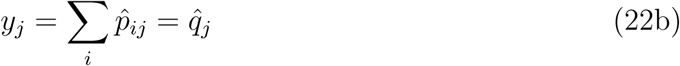

and

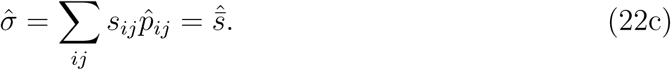

We can write

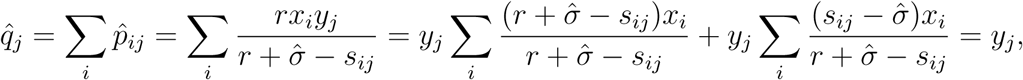

since **x** ∈ Δ_*m*_ and 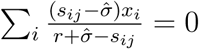 and by (15c).

An analogous computation shows that 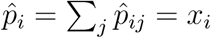. This implies

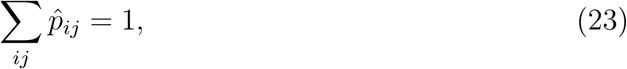

because **x** ∈ Δ_*m*_.

We can rewrite (21) as

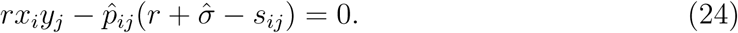

Summing (24) over all *i* and *j*, yields

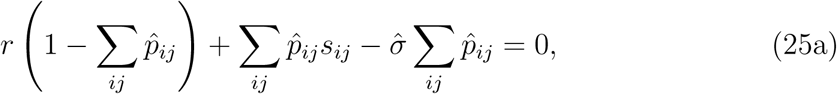

which implies

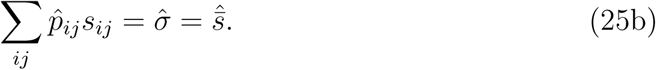

Now, we can write (24) as

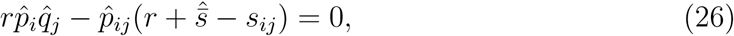

which is (2). □

With this characterization of the polymorphism at hand, we can prove Theorem 3.

*Proof of Theorem 3.* If 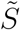 is such that there exist no vectors *x, y* and values *r* and 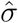 such that all conditions in (15) are fulfilled, then no equilibrium exists. The following argument shows that if there is a solution that satisfies (15), then there are infinitely many provided *m* ≠ *n*.

Assume *m* < *n* and suppose there exists **x** ∈ Δ_*m*_, **y** ∈ Δ_*n*_ and 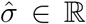 such that holds. Then 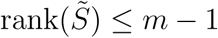 which, because of the rank-nullity theorem, implies that 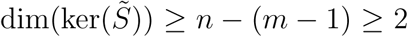. This means that at least one additional linearly independent vector **y**′ exists in the kernel of 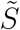. This solution vector does not necessarily lie in the simplex. However, 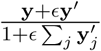 defines a one dimensional manifold that lies in the simplex for 0 < *ϵ* < *ϵ*^*^ with *ϵ*^*^ > 0 sufficiently small and is a solution of (15b). Therefore, 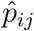 is at least one-dimensional. □

The characterization of internal equilibria given by Lemma 2 yields necessary conditions for the existence of equilibria with an equal number of alleles present.

#### Proposition 2.

*If n* = *m and an isolated equilibrium* 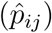 *of* (2) *satisfying* (13) *exists, then the following holds:*

a. *No row or column of* 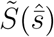 *consists only of entries of the same sign.*
b. 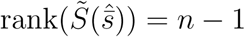.
c. 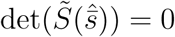.
d. *There is no other equilibrium with the same* 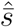.

*Proof.* Let **y** be the vector for which (15b) holds. First, assume that there exists a row *i* of 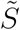 with all entries positive. This implies 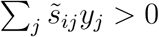, a contradiction, because (15b) holds for **y**. An analogous argument works for column *j* and **x**. This yields statement (a).

The rank of 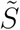 has to be smaller than *n*, because (15b) can only have a nontrivial solution if 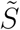 is singular.

Now, assume that 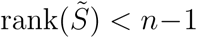. Then by the rank-nullity theorem, 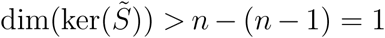. Therefore, at least one additional linearly independent vector **y**′ exists in the kernel of 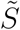. This solution vector does not necessarily lie in the simplex. However, 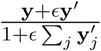 defines a one dimensional manifold that lies in the simplex for 0 < *ϵ* < *ϵ*^*^ with *ϵ*^*^ > 0 sufficiently small and is a solution of (15b). This contradicts the assumption of an isolated equilibrium. Thus statement (b) is true. Statement (c) immediately follows. □

If 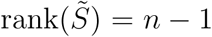, then **y** spans the kernel of 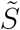. Hence no other equilibrium with the same 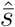 is possible. Hence, statement (d) follows. □

Note, that statements (a)-(d) are not sufficient conditions for the existence of an internal equilibrium. In particular, statement (a) of Proposition 2 does not imply that a positive solution vector exists. It is also not clear if there always exists a 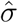 simultaneously fulfills (15a) and statements (a) and (c) of Proposition 2.

In simple situations Lemma 2 and Proposition 2 allow us to get further results.

Statements (c) and (d) combined yield an upper bound for the number of internal equilibria, since each zero of 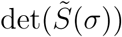 gives rise to at most one equilibrium. Since 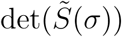 is a rational function of *σ*, the degree of the numerator polynomial determines the maximal possible number of internal equilibria. For small numbers of alleles this argument gives rise to a feasible method to determine all admissible internal equilibria. In fact, for two alleles we show in the following that there is at most one internal equilibrium, which is also true for three alleles with a centrosymmetric fitness scheme (see SI).

If *S*′ is the scaled fitness scheme with respect to the equilibrium *E*, then

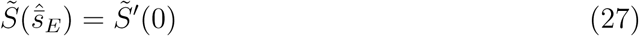

holds, by Remark 1. Therefore, statement (c) of Proposition 2 implies det 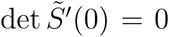. This yields an additional identity that the scaled fitnesses 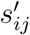 have to satisfy. For small numbers of alleles, this can be solved explicitly and implicit coordinates for *E* can be obtained.

### Two explicit cases with an equal number of alleles at both loci

#### Two alleles

In the two allele case the following Theorem is already known from Bank, Bürger and Hermisson (2012), but we present a different proof.

##### Theorem 4.

*System* (1) *restricted to two alleles at each locus has a unique internal equilibrium if and only if there exists* 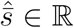 *such that* 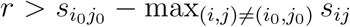 *and* 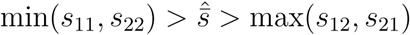 *or the reverse order holds. Here*, 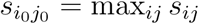. *If the equilibrium exists, it is unstable.*

*Proof.* (⇒) Without loss of generality, we assume 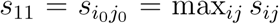. Since the equilibrium is unique, statement (a) of Proposition 2 yields

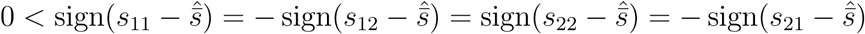

This implies

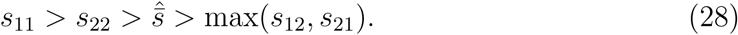

Additionaly, Lemma 2 tells us that (15a) has to hold, which we combine with (27) to conclude

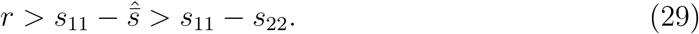

(⇐) Without loss of generality, we assume 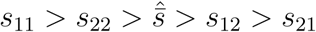. By applying Proposition 1, we know that the monomorphism *A*_1_*B*_1_ is asymptotically stable for all *r*, while the monomorphisms *A*_1_*B*_2_ and *A*_2_*B*_1_ are unstable for all *r*. The monomorphism *A*_2_*B*_2_ is asymptotically stable if *r* > *s*_11_ − *s*_22_. This is exactly the condition 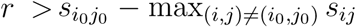 with (*i*_0_, *j*_0_) = (1, 1). There are no other boundary equilibria, since by Theorem 3 there is no equilibrium on the edges where one allele is fixed. Thus, the sum of all indices on the boundary is 2. Since the index theorem by Hofbauer (1990) implies that the sum over the indices of all saturated equilibria is 1, the sum of all indices of internal equilibria has to equal −1 (they are saturated by definition). This entails an odd number of internal equilibria, because the index of a hyperbolic equilibrium is either +1 or −1.

However, the degree of the numerator polynomial (in *σ*) of the determinant of the corresponding matrix 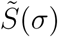 is two (see SI for the exact expression). Hence, there are at most two values of *σ* such that 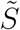 is singular and this is a necessary condition by statement (b) of Proposition 2. Thus there are up to two internal equilibria. By the index argument above, only an odd number of internal equilibria is possible and therefore, there is exactly one internal equilibrium. It has index −1, which implies an odd number of positive eigenvalues and subsequently its instability. □

With an additive scaling of the fitness matrix, detailed in Remark 1, we can easily express the equilibrium coordinates of the internal equilibrium.

##### Corollary 3.

*Let S be the scaled fitness scheme such that* 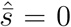 *at the internal equilibrium and s*_22_ *= š*_22_ *holds, with*

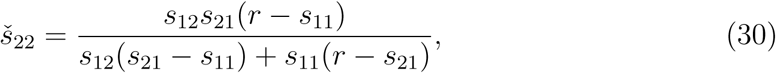

*then the equilibrium is given by*

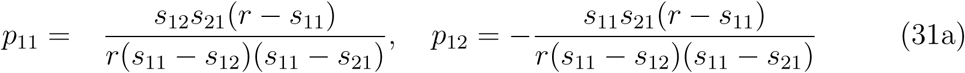

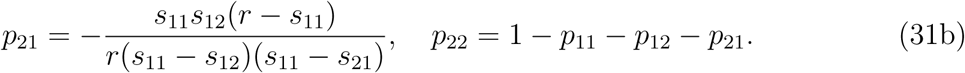

*It is admissible if* sign(*s*_11_*s*_22_) = sign(*s*_21_*š*_22_) = −sign(*s*_11_*š*_22_) = −1 *and*

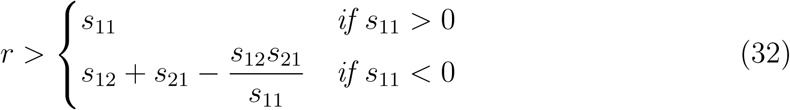

*Proof.* According to Remark 1, we can formally set 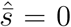 in 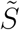, restricted to two alleles per locus, which simplifies further analysis. 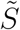 is a singular matrix if *s*_22_ *= š*_22_. Subsequently we solve (15b) and (15c) and use formula (16) to derive (30). Admissibility conditions follow from Theorem 4, after one resolves the dependence of *š*_22_ on *r* with repect to sign(*š*_22_) and *r* > *š*_22_. See also SI. □

#### Three alleles

We now investigate the case of three alleles at each locus. Additionally, we assume a centrosymmetric fitness scheme:

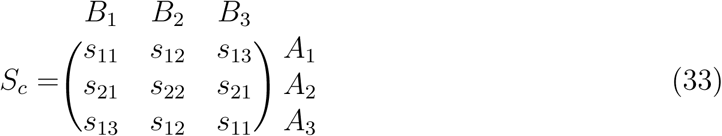

In the sense that the matrix is the same but the entries correspond to haplotypes formed by single alleles rather than by diploid genotypes, *S*_*c*_ is formally equivalent to the well-studied symmetric viability model. Similarly, it implies that the fitnesses stay the same under a simultaneous exchange of alleles *A*_1_ with *A*_3_ and *B*_1_ with *B*_3_. Unfortunately, it violates assumption (ℋ), since *s*_32_ = *s*_12_ and *s*_23_ = *s*_21_ and thus every point on the edges connecting the monomorphisms *A*_1_*B*_2_ with *A*_3_*B*_2_ and *A*_2_*B*_1_ with *A*_2_*B*_3_ is an equilibrium by Lemma 1.

Since we are mainly interested in the stability of a potential internal equilibrium, from now on, we assume that an internal equilibrium exists with mean fitness at this equilibrium given by 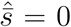 (see Remark 1).

This simplifies (14) considerably, because

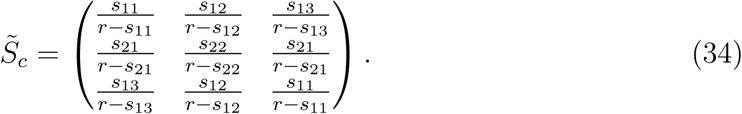

The coordinates of the equilibrium for which *S* is scaled, are described by the following

##### Proposition 3.

*Let* 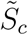 *be as in* (33) *and s*_11_ ≠ *s*_13_. *In addition, define*

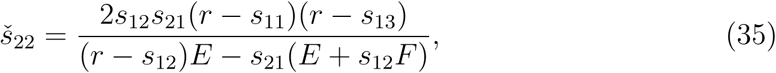

*with E* = *s*_11_(*r* − *s*_13_) + *s*_13_(*r* − *s*_11_) *and F* = *s*_11_ + *s*_13_ − 2*r.*

*Then, an equilibrium, given by* (16) *satisfying* (13), *exists if and only if*

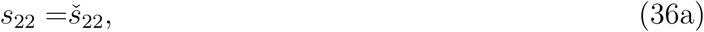

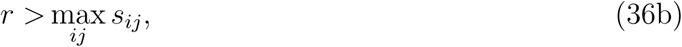

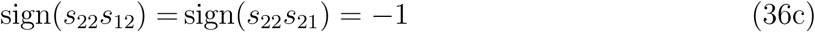

*and*

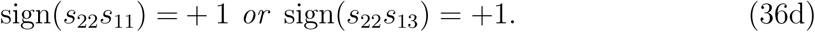

*Its coordinates are*

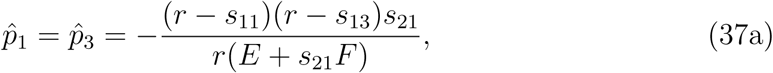

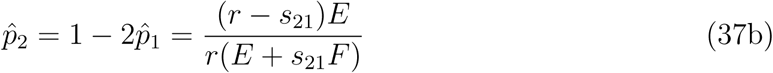

*and*

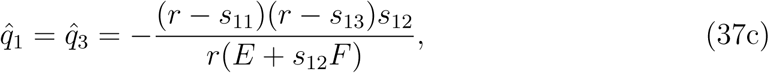

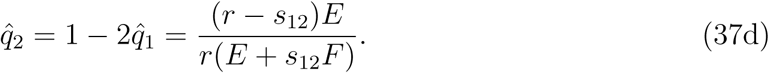

*Proof.* (⇒) If the equilibrium exists, we can scale *S* such that 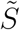 is given by (33) and 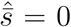. Then, condition (15a) simplifies to

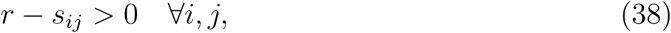

which implies (35b). The equation in statement (c) of Proposition 2 for (33) is linear in *s*_22_ and thus has the unique solution *š*_22_. Statemant (a) implies (35c) and (35d), because otherwise, at least one row or column of (33) consists only of entries of the same sign.

(⇐) The determinant of (33) is linear in *s*_22_ and is zero if *s*_22_ *= š*_22_. Thus, 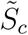 is a singular matrix and (15b) can have a admissible solution. Equating the first and third row of (15b) yields

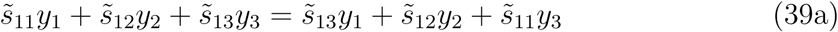

which simplifies to

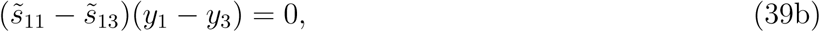

where 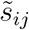 denote the entries of 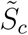 Since *s*_11_ ≠ *s*_13_, (38b) holds if and only if *y*_1_ = *y*_3_. By applying ∑_*j*_ *y*_*j*_ = 1, we can express (15b) in terms of *y*_1_ and solve for it. An analogous argument yields *x*_1_ and we get (36) (at equilibrium 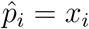 and 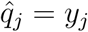).

By Lemma 2 these coordinates determine an equilibrium for (1). However, it remains to show that 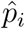 and 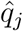 given by (36) are positive, given the conditions (35). Without loss of generality, *s*_22_, *š*_22_ > 0. This implies that *s*_12_, *s*_21_ < 0. Let *h*_1_ denote the numerator and *h*_2_ the denominator of *š*_22_ given in (34). Since *h*_1_ is clearly positive, the same has to hold for *h*_2_. Condition (37) also holds for *š*_22_ and simplifies to

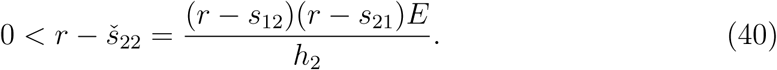

Therefore, we conclude that sign(*E*) = sign(*h*_2_) = sign(*s*_22_) = 1, since the first two factors of the numerator are positive because of (37). This is possible only if (35d) is true, since otherwise, *s*_11_, *s*_13_ < 0 imply *E* < 0, provided (37) holds. Because of (37), *F* < 0 and thus the denominators of (36) are all positive. A simple check reveals that this also holds for each numerator of (36).

For any fitness scheme that can be achieved by adding a constant to *s*_*ij*_, an equilibrium exists and its coordinates are given by (36) where a constant is added to each *s*_*ij*_. □

If *s*_11_ = *s*_13_, then it is clear that 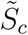 given by (33) is singular. However, if 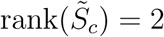, then one can easily check that the equilibrium is not admissible. If 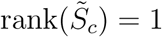, then the equilibrium is not isolated by Proposition 2.

##### Corollary 4.

*The equilibrium given by* (36) *is centrosymmetric, i.e.*,

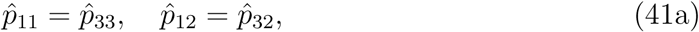

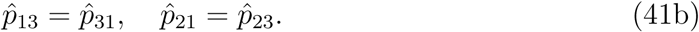

*Proof.* Using (16) one can derive the exact expressions for 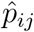 given (36), they are rather lengthy and thus only shown in the SI. Then it is easy to check that they satisfy (40). □

Before we prove the instability of an internal poylmorphism, if it exists, we note that Corollary 4 gives rise to the coordinate transformation (𝒰) inspired by Feldman and Karlin (1970), which provides the starting point for the proof.

We define:

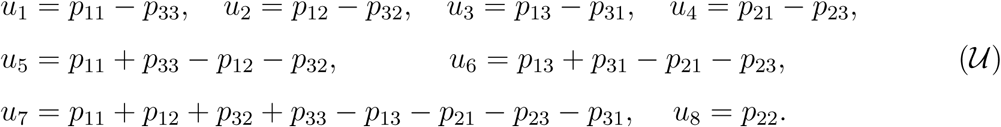

From (𝒰) and the simplex condition ∑_*ij*_*p*_*ij*_ = 1, we derive the reverse transformation:

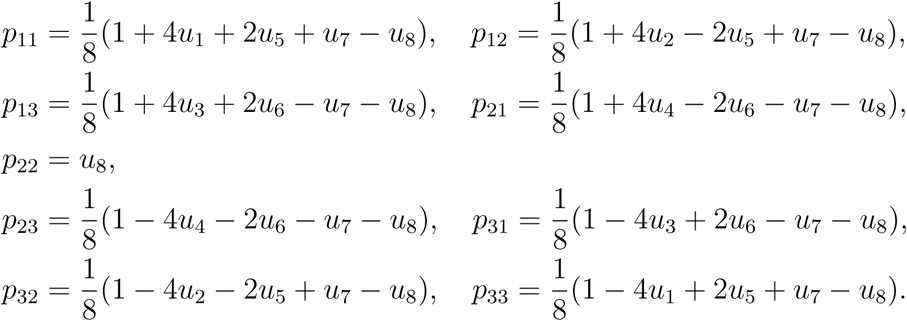

With this, we now derive the transformed system of equations from (1). However, this rather lengthy system of ODEs is only shown in the SI. The equilibrium coordinates 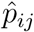 are also transformed into the equilibrium *û*_*i*_ (see SI). Because of Corollary 4 and (𝒰), *û* = 0 for *i* = 1, 2, 3 and 4.

With these prerequisites, we can prove the main theorem of this section.

##### Theorem 5.

*If the polymorphic equilibrium* (36) *of* (1) *under the fitness scheme S*_*c*_*(eq. 32) exists, then it is unstable.*

*Proof.* For the new system in (*u*_*i*_), we compute the Jacobian *J*_*u*_ (see SI for the derivation). We evaluate it at the hyperplane *H* given by 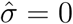 and *û*_*i*_ = 0 for *i* = 1, 2, 3 and 4. Clearly, *H* contains the equilibrium.

The resulting 8 × 8 matrix is then in block diagonal form

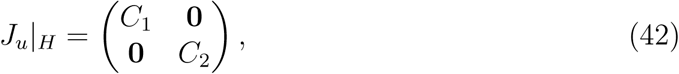

where each block is a square matrix of dimension 4. As the determinant of the full matrix is the product of the determinants of *C*_1_ and *C*_2_, we can analyze them separately.

The resulting expression for det(*C*_1_) (see SI for the expression) is then evaluated at the coordinates *û*_*i*_, *i* = 5, 6, 7 and 8. After some simplification, it can be written as:

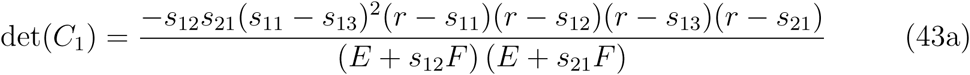

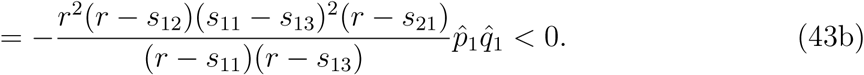

This inequality holds, because condition (37) ensures that the ratio in inequality (42b) is positive and 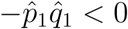, since (36) is an admissible equilibrium.

The determinant of *C*_1_ is the product of the four eigenvalues and we can thus conclude that at least one of them has to be positive. This implies that the equilibrium is unstable. □

##### Remark 4.

*Neither Proposition 3 nor Theorem 5 states that the equilibrium it concerns is the only internal equilibrium. If there were two isolated equilibria E*_1_ *and E*_2_ *with* 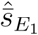 and 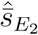 *respectively then both Proposition 3 and Theorem 5 would remain valid if we set once* 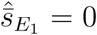 *and once* 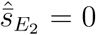. *However, in the SI we prove uniqueness of the internal equilibrium, if conditions* (35c) *and* (35d) *of Proposition 3 hold.*

## Discussion

We have conducted a rather general mathematical analysis of haploid two-locus multiallele dynamics with constant selection and recombination. The model we use is the standard continuous-time model for selection on haploids with recombination, see eq. (1).

In the first section, we provide conditions for stability of hyperbolic monomorphisms and show that at least one of them is always stable. If both loci exhibit the same number of alleles, then there are at most *n* stable monomorphisms. If the number of alleles is unequal, then the smaller of them is the upper bound for the number of stable monomorphism. We also characterize the fitness matrices such that the only stable monomorphism is the same for every *r* > 0 and claim that it is also globally asymptotically stable for all *r*. But this remains unproven.

However, we use ideas about dominating strategies from game theory to prove global stability in section 2. If for a fixed background allele the haplotype formed with one allele is greater than with another allele and this holds for every background allele, than the fitter allele is the dominating allele. The dominated allele goes extinct. Subsequently, if one allele dominates all other alleles on the same locus, then they all go extinct and the dominating allele is fixed. Thus the two-locus multi-allele problem is reduced to a one-locus multi-allele problem. There, it is known that the allele with the highest fitness gets fixed. Similar to the game theoretic problem, we apply a quasi-concave Lyapunov function to prove global convergence. Speaking informally, this approach helps to get control over the terms of linkage disequilibria. These terms are introduced when we consider population genetic dynamics with more than one locus and they are the reason why the usual Lyapunov methods that work for one-locus models break down. Potentially, other multi-locus convergence problems could be treated by means of quasi-concave Lyapunov functions.

Lemma 2 represents a very useful and intuitive characterization of polymorphisms in terms of two linear homogeneous systems of equations. Solvability of both systems in (15) is necessary and sufficient for the existence of internal equilibria. If the numbers of alleles at the two loci are different, then one of the systems is overdetermined and has, in general, no solution. However, in the degenerate case, where a solution exists, we showed that there is a manifold of solutions. This means for an unequal number of alleles at the loci, there is either no internal equilibrium or there are infinitely many. This immediate consequence is formalized in Theorem 3. If the two loci have the same number of alleles, we state necessary conditions for the existence of an isolated internal equilibrium in Proposition 2.

If there are either two or three alleles (with centrosymmetric fitnesses) at both loci, these general results on the existence of internal equilibria are used to establish uniqueness of the polymorphism and its instability if it exists.

With two alleles at each locus it is rather straightforward to prove uniqueness of the internal equilibrium by combining an index theorem by Hofbauer (1990) with Proposition 1, Theorem 3 and Proposition 2. The index theorem also entails that the equilibrium is a saddle point. This approach shows the uniqueness and instability of the polymorphism simultaeously and is not as technical and computationally difficult as that of Bank, Bürger and Hermisson (2012).

For three alleles at both loci, we need additional assumptions to establish an analogous result. We assume a centrosymmetric fitness scheme, generalizing that of the classical symmetric viability two-locus two-allele diploid model.

After scaling *S*_*c*_ such that 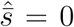, we determine the exact equilibrium coordinates, which also exhibit centrosymmetry. This allows us to apply a coordinate transformation that exploits this symmetry. With its help, instability of the internal equilibrium is shown. Note that in this proof we do not need the uniqueness of the equilibrium. However, in the SI, we show that the internal equilibrium for this centrosymmetric three-allele model is in fact unique.

Conditions (35) together with Proposition 1 imply that three monomorphisms are locally asymptotically stable. This in turn implies, by Theorem 4, that in each of the three two-allele subsystems spanned by these three monomorphisms a unique biallelic unstable polymorphism exists. Generalizing this argument in a rather speculative fashion, we claim that 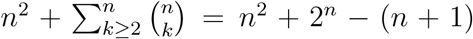 is the maximum number of isolated equilibria for system (1) with *n* alleles at both loci. There, we assume that for *k* > 1 alleles at both loci 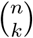 isolated equilibria exist. This is only proven for *n* = 2, because the centrosymmetry assumption in our treatment of *n* = 3 entails that four monomorphisms are not isolated, since Lemma 1 ensures the existence of edges where every point is an equilibrium. However, if we set *s*_23_ = *s*_21_ + *ϵ* and *s*_32_ = *s*_12_ + *ϵ*, then local perturbation theory implies that there are 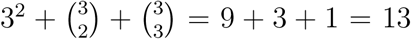 isolated equilibria for *ϵ* sufficiently small. According to the claim we made above, this should hold for the general three allele case. The claim also implies uniqueness of the equilibrium with all *n* alleles present.

With the results presented here and the assumption that the trajectories converge to equilibrium points, it is clear that genetic variation, if it is maintained at two loci through haploid selection and recombination, only occurs with the same number (larger or equal 3) of alleles at both loci. If at one locus exactly two alleles occur or exactly three, which in addition are centrosymmetric, then genetic variation is always lost regardless of the number of alleles at the other locus. Ultimately, the population is fixed for one allele at each locus. Additionally, variation vanishes if the fitness scheme is of the form given in Theorem 2. The genotype with the maximal fitness becomes fixed.

## Supporting information

Supplement Mathematica code as pdf

## Acknowledgements

We thank Reinhard Bürger for useful discussions and Josef Hofbauer for pointing to a very similar problem about global stability in game theory and help with the formulation and proof of Lemma 2.

Financial support by the Austrian Science Fund (FWF) through the Vienna Graduate School of Population Genetics (Grant W1225) to MP is gratefully acknowledged. This work emerged from a visit of MP to MWF in 2018 made possible by a scholarship from the Austrian Marshall Plan Foundation.

## References

Bank C., Bürger R., Hermisson J. 2012. Limits to parapatric speciation: Dobzhansky-Muller incompatibilities in a continent-island model. Genetics, 191, 845–863.

Feldman M.W. 1971. Equilibrium studies of two locus haploid populations with recombination. Theor. Popul. Biol. 2, 299–318.

Feldman M.W., Karlin S. 1970. Linkage and selection: Two locus symmetric viability model. Theor. Popul. Biol. 1, 39–71.

Hofbauer J. and Su J.J. 2016. Global stability of spatially homogeneous equilibria in migration-selection models. SIAM J. Appl. Math. 76, 578–597.

Hofbauer J. 1990. An index theorem for dissipative semiflows. Rocky Mountain Journal of Mathematics. 20, 1017–1031.

Kirzhner V.M. and Lyubich Y. 1997. Multilocus dynamics under haploid selection. J. Math. Biol. 35: 391–408

Nagylaki T. 1992. Introduction to Theoretical Population Genetics. Springer-Verlag

Novak S., Barton N.H. 2017. When does frequency-independent selection maintain genetic variation? Genetics 207: 653–668

Rutschman D. 1994. Dynamics of the two-locus haploid model. Theor. Popul. Biol. 45, 167–176.

