## Supplement Mathematica code as pdf for "Loss of genetic variation in the two-locus multiallelic haploid model"

### 3 alleles at both loci:

#### Uniqueness:

To prove uniqueness we scale the fitness-matrix such that s[2,2]=0.

We want to show that there is at most one $\bar{s}$ for which det($\tilde{S}$)=0 with $r+\bar{s}-s[i,\ j]>0$. And recall Lemma 2 from the main text.

```
Stildecentro = S̃[3] /. centrosym[3] /. s[2, 2] → 0;
MatrixForm[Stildecentro]
```

Matritzenform

$$\begin{pmatrix} \frac{-\bar{s}+s[1,1]}{r+\bar{s}-s[1,1]} & \frac{-\bar{s}+s[1,2]}{r+\bar{s}-s[1,2]} & \frac{-\bar{s}+s[1,3]}{r+\bar{s}-s[1,3]} \\ \frac{-\bar{s}+s[2,1]}{r+\bar{s}-s[2,1]} & -\frac{\bar{s}}{r+\bar{s}} & \frac{-\bar{s}+s[2,1]}{r+\bar{s}-s[2,1]} \\ \frac{-\bar{s}+s[1,3]}{r+\bar{s}-s[1,3]} & \frac{-\bar{s}+s[1,2]}{r+\bar{s}-s[1,2]} & \frac{-\bar{s}+s[1,1]}{r+\bar{s}-s[1,1]} \end{pmatrix}$$

```
detStc = Simplify[Det[Stildecentro]]
```

vereinfache Determinante

$$-\Big(\Big(r^2\,(s[1,1]-s[1,3])\,\Big(\bar{s}^3\,(s[1,1]-2\,s[1,2]+s[1,3]-2\,s[2,1])+2\,(r-s[1,1])\,s[1,2]\,(r-s[1,3])\,s[2,1]+\bar{s}^2\,(s[1,2]\,s[1,3]+2\,r\,(s[1,1]-2\,s[1,2]+s[1,3]-2\,s[2,1])+4\,s[1,2]\,s[2,1]+s[1,3]\,s[2,1]+s[1,1]\,(s[1,2]-2\,s[1,3]+s[2,1]))+\bar{s}\,\big(r^2\,(s[1,1]-2\,s[1,2]+s[1,3]-2\,s[2,1])-3\,s[1,2]\,(s[1,1]+s[1,3])\,s[2,1]+r\,(s[1,3]\,s[2,1]+s[1,1]\,(s[1,2]-2\,s[1,3]+s[2,1])+s[1,2]\,(s[1,3]+6\,s[2,1]))\big)\Big)\Big)\Big/\Big((r+\bar{s})\,(r+\bar{s}-s[1,1])^2\,(r+\bar{s}-s[1,2])\,(r+\bar{s}-s[1,3])^2\,(r+\bar{s}-s[2,1])\Big)\Big)$$

##### Total number of possible orderings:

We need a good control over the sign of the determinatn at various points. There it proves helpful to count the possible orderings of the 5 fitness parameters.

There are 5!=120 in total. However, since the number of equilibria is not changed if allele A1 is renamed A3 and locus A exchanges name with locus B, we can basically identify s[1,2] with s[2,1] and s[1,3] with s[1,1] in the orderings: (It can help to think of this process as choosing balls from a urn. Two are blue (s[1,2] and s[2,1]), two are green (s[1,1] and s[1,3]) and one is black (s[2,2]). How many orderings are there without refilling the urn? 30)

```
redpermut = Tally[Permutations[{s[1, 1], s[1, 2], s[1, 1], s[1, 2], s[2, 2]}]];
```

zähle··· Permutationen

```
Length[redpermut]
```

Länge

30

```
colorballpermut = Tally[Permutations[{Green , Green, Blue, Blue, Black}]];
                        zähle…   Permutationen    grün     grün    blau  blau  schwarz
Length[colorballpermut]
Länge
```

30

Lemma 2 of the main text implies that $s[2, 2] - \bar{s}$ and $s[1, 2] - \bar{s}$ have to have opposite signs. Similarly $s[2, 1] - \bar{s}$ has to have the same sign as $s[1, 2] - \bar{s}$. Thus we can exclude all cases from above where s[1,2] occurs above and below s[2,2]. Also we need at least three entries of $\tilde{S}$ have the same sign, therefore, if s[2,2] is the minimum of an ordering, then s[1,2] can not be the direct upper bound. See also conditions (36c) and (36d) in the main text.

Somehow different proof-methods work better conditioned on maximality or minimality of s[2,2]:
If s[2,2] is max or close to the top, it works well to set s[2,2] to zero and make a coordinate transformation of the determinant such that the interval (-r,0) maps to the positive halfaxis. Using Descarte sign rule, we conclude that at most one zero can exist in this interval and this is thus true for the number of internal equilibria.
The relevant cases are:
s[2,2]>s[1,1]>s[1,1]>s[1,2]>s[1,2]
s[2,2]>s[1,1]>s[1,2]>s[1,1]>s[1,2]
s[2,2]>s[1,1]>s[1,2]>s[1,2]>s[1,1]

s[1,1]>s[2,2]>s[1,1]>s[1,2]>s[1,2]
s[1,1]>s[2,2]>s[1,2]>s[1,1]>s[1,2]
s[1,1]>s[2,2]>s[1,2]>s[1,2]>s[1,1]

s[1,1]>s[1,1]>s[2,2]>s[1,2]>s[1,2]

If s[2,2] is minimal, or close to the minimum, then the interval were admissible $\bar{s}$ exist is not the same as above. However, there are five poles which define 6 intervalls in which the determinant is continuous. Thus we can compute the sign at the start and endpoints of these intervals. If the sign changes within one interval, we can follow by middle value theorem that there has to be an odd number of zeroes in between. We find always three intervals conanting at least one zero and since the degree is three there are at most three zeroes in total.
The cases analysed here are:
s[2,2]<s[1,1]<s[1,1]<s[1,2]<s[1,2]
s[2,2]<s[1,1]<s[1,2]<s[1,1]<s[1,2]
s[2,2]<s[1,1]<s[1,2]<s[1,2]<s[1,1]

s[1,1]<s[2,2]<s[1,1]<s[1,2]<s[1,2]
s[1,1]<s[2,2]<s[1,2]<s[1,1]<s[1,2]
s[1,1]<s[2,2]<s[1,2]<s[1,2]<s[1,1]

s[1,1]<s[1,1]<s[2,2]<s[1,2]<s[1,2]

The proofs are simplified by setting the smaller s[1,1] and s[1,2] in the above orderings to s[1,3] and s[2,1] respectively.

### Interval mapping method:

#### s[2,2] max, then s[1,1]. Internal equilibrium only possible if s[1,2], s[2, 1] < $\bar{s}$.

```
assparas22 =
 FullSimplify[Reduce[{r > -s[1, 1] > 0 ≥ s[2, 2] > s[1, 1] > s[2, 1], s[1, 1] > s[1, 2],
 vereinfache voll··· reduziere
     s[1, 1] > s[1, 3]} /. s[2, 2] → 0]]
```

```
s[2, 1] < s[1, 1] < 0 && s[1, 3] < s[1, 1] && s[1, 2] < s[1, 1] && r + s[1, 1] > 0
```

Since $s[1, 1] > \bar{s}$, $0 < r + \bar{s}$-s[2,2]<r+s[1,1].

The admissible $\bar{s}$ has to be between -r and 0. We use the transformation $z = -(\bar{s} + r)/\bar{s}$ to map this interval to the positive axis.

```
Solve[z == -(s̄ + r) / s̄, s̄]
löse
```

$$\left\{\left\{\bar{s} \to -\frac{r}{1+z}\right\}\right\}$$

```
detposz =
 Collect[Together[Numerator[detStc] (1 + z)^3 /. s̄ → -r / (1 + z)], z, FullSimplify]
 gruppier··· zusammen  Zähler                                          vereinfache vollstär
```

```
-2 r^2 z^3 (r - s[1, 1]) s[1, 2] (r - s[1, 3]) (s[1, 1] - s[1, 3]) s[2, 1] -
 r^2 s[1, 2] (s[1, 1] - s[1, 3]) (r s[1, 1] + (r + 2 s[1, 1]) s[1, 3]) s[2, 1] +
 r^2 z (s[1, 1] - s[1, 3]) (-6 s[1, 1] s[1, 2] s[1, 3] s[2, 1] + r^2 (s[1, 2] s[1, 3] +
        (2 s[1, 2] + s[1, 3]) s[2, 1] + s[1, 1] (s[1, 2] - 2 s[1, 3] + s[2, 1]))) +
 r^2 z^2 (s[1, 1] - s[1, 3]) (r^3 (s[1, 1] - 2 s[1, 2] + s[1, 3] - 2 s[2, 1]) -
    6 s[1, 1] s[1, 2] s[1, 3] s[2, 1] + 3 r s[1, 2] (s[1, 1] + s[1, 3]) s[2, 1] +
    r^2 (s[1, 3] (s[1, 2] + s[2, 1]) + s[1, 1] (s[1, 2] - 2 s[1, 3] + s[2, 1])))
```

We can write detposz as c0+c1z+c2z^2+c3z^3:

```
{c0, c1, c2, c3} = CoefficientList[detposz, z]
                   Liste der Koeffizienten
```

```
{-r^2 s[1, 2] (s[1, 1] - s[1, 3]) (r s[1, 1] + (r + 2 s[1, 1]) s[1, 3]) s[2, 1],
 r^2 (s[1, 1] - s[1, 3]) (-6 s[1, 1] s[1, 2] s[1, 3] s[2, 1] + r^2 (s[1, 2] s[1, 3] +
        (2 s[1, 2] + s[1, 3]) s[2, 1] + s[1, 1] (s[1, 2] - 2 s[1, 3] + s[2, 1]))),
 r^2 (s[1, 1] - s[1, 3]) (r^3 (s[1, 1] - 2 s[1, 2] + s[1, 3] - 2 s[2, 1]) -
    6 s[1, 1] s[1, 2] s[1, 3] s[2, 1] + 3 r s[1, 2] (s[1, 1] + s[1, 3]) s[2, 1] +
    r^2 (s[1, 3] (s[1, 2] + s[2, 1]) + s[1, 1] (s[1, 2] - 2 s[1, 3] + s[2, 1]))),
 -2 r^2 (r - s[1, 1]) s[1, 2] (r - s[1, 3]) (s[1, 1] - s[1, 3]) s[2, 1]}
```

```
FullSimplify[c1 < 0 /. r → -s[1, 1], assparas22]
vereinfache vollständig
```

```
True
```

```
FullSimplify[c2 < 0 /. r → -s[1, 1], assparas22]
vereinfache vollständig
```

```
True
```

```
FullSimplify[c2 - c1 < 0 /. r → -s[1, 1], assparas22]
vereinfache vollständig
```

```
True
```

```
FullSimplify[c3 < 0, Flatten[{assparas22}]]
vereinfache vollständig    ebne ein
Reduce[Flatten[{c1 < 0 < c2, assparas22}]]
reduzi···  ebne ein
```

```
True
```

```
False
```

```
Eplus = r (s[1, 1] + s[1, 3]) + 2 s[1, 1] s[1, 3];
```

```
FullSimplify[c0 > 0, Flatten[{assparas22, Eplus < 0}]]
vereinfache vollständig    ebne ein
FullSimplify[c0 < 0, Flatten[{assparas22, Eplus > 0}]]
vereinfache vollständig    ebne ein
```

```
True
```

```
True
```

If Eplus<0, then there can be the sign vectors of the coefficents:
{+,-,-,-}, {+,+,-,-}, {+,+,+,-}.
With Descartes we conclude that there can be at most one zero in the relevant interval.

```
Simplify[Reduce[Flatten[{c1 < 0, assparas22, Eplus > 0}]],
vereinfache reduzi··· ebne ein
  Flatten[{assparas22, Eplus > 0}]]
  ebne ein
Simplify[Reduce[Flatten[{c2 < 0, assparas22, Eplus > 0}]],
vereinfache reduzi··· ebne ein
  Flatten[{assparas22, Eplus > 0}]]
  ebne ein
```

```
True
```

```
True
```

If Eplus>0, then the sign vector of the coefficients is {-,-,-,-}. There is no zero in the relevant interval.

Eplus > 0 is in this case equivalent to r<$-\frac{2\,s[1,1]\,s[1,3]}{s[1,1]+s[1,3]}$. So if r is too small, then there cannot be an internal equilibrium with this general order.

#### s[1,1] max, then s[2,2] or s[1,3]. Internal equilibrium only possible if s[1, 2], s[2, 1] < s̄.

```
assparas11 = FullSimplify[Reduce[{s[1, 1] > s[2, 2] > s[1, 2],
             vereinfache voll··· reduziere
      s[1, 1] > s[1, 3], s[2, 2] > s[2, 1], r > s[1, 1]} /. s[2, 2] → 0]]
```

```
s[1, 2] < 0 && s[2, 1] < 0 && r > s[1, 1] &&
 (s[1, 1] > 0 || s[1, 3] > 0) && (s[1, 1] > s[1, 3] || s[1, 3] ≤ 0)
```

0 < r + s̄ − s[1, 1] < r − s[1, 1] , because s̄ < 0.

Again, s̄ < 0 and greater than the first pole -r+s[1,1]. We map this interval to the positive halfaxis for the numerator of the determinant:

```
Solve[z == (-s̄ - r + s[1, 1]) / s̄, s̄]
löse
```

$$\left\{\left\{\bar{s} \to \frac{-r + s[1, 1]}{1 + z}\right\}\right\}$$

```
det0pole1 =
 Collect[Together[Numerator[detStc] (1 + z)^3 /. s̄ → (-r + s[1, 1])/(1 + z)], z, FullSimplify]
 gruppier··· zusammen  Zähler                                                   vereinfache vollstär
```

```
r^2 (r - s[1, 1]) (s[1, 1] - s[1, 2]) (s[1, 1] - s[1, 3])^2 (s[1, 1] - s[2, 1]) -
 2 r^2 z^3 (r - s[1, 1]) s[1, 2] (r - s[1, 3]) (s[1, 1] - s[1, 3]) s[2, 1] +
 r^2 z^2 (r - s[1, 1]) (s[1, 1] - s[1, 3])
  (r^2 (s[1, 1] - 2 s[1, 2] + s[1, 3] - 2 s[2, 1]) + 3 s[1, 2] (-s[1, 1] + s[1, 3]) s[2, 1] +
    r (s[1, 3] (s[1, 2] + s[2, 1]) + s[1, 1] (s[1, 2] - 2 s[1, 3] + s[2, 1]))) +
 r^2 z (r - s[1, 1]) (s[1, 1] - s[1, 3]) (s[1, 1] (s[1, 2] (s[1, 3] - 2 s[2, 1]) +
         s[1, 3] s[2, 1] + s[1, 1] (s[1, 2] - 2 s[1, 3] + s[2, 1])) + r (2 s[1, 1]^2 +
         s[1, 3] s[2, 1] - 3 s[1, 1] (s[1, 2] + s[2, 1]) + s[1, 2] (s[1, 3] + 2 s[2, 1])))
```

It is a polynomial in z that is written as d0+d1z+d2z^2+d3z^3:

```
{d0, d1, d2, d3} = CoefficientList[det0pole1, z]
                   Liste der Koeffizienten
```

```
{r^2 (r - s[1, 1]) (s[1, 1] - s[1, 2]) (s[1, 1] - s[1, 3])^2 (s[1, 1] - s[2, 1]),
 r^2 (r - s[1, 1]) (s[1, 1] - s[1, 3]) (s[1, 1] (s[1, 2] (s[1, 3] - 2 s[2, 1]) +
         s[1, 3] s[2, 1] + s[1, 1] (s[1, 2] - 2 s[1, 3] + s[2, 1])) + r (2 s[1, 1]^2 +
         s[1, 3] s[2, 1] - 3 s[1, 1] (s[1, 2] + s[2, 1]) + s[1, 2] (s[1, 3] + 2 s[2, 1]))),
 r^2 (r - s[1, 1]) (s[1, 1] - s[1, 3]) (r^2 (s[1, 1] - 2 s[1, 2] + s[1, 3] - 2 s[2, 1]) +
    3 s[1, 2] (-s[1, 1] + s[1, 3]) s[2, 1] +
    r (s[1, 3] (s[1, 2] + s[2, 1]) + s[1, 1] (s[1, 2] - 2 s[1, 3] + s[2, 1]))),
 -2 r^2 (r - s[1, 1]) s[1, 2] (r - s[1, 3]) (s[1, 1] - s[1, 3]) s[2, 1]}
```

```
FullSimplify[d0 > 0, assparas11]
```
vereinfache vollständig

```
FullSimplify[d3 < 0, assparas11]
```
vereinfache vollständig

```
Simplify[Reduce[Flatten[{d1 > 0, assparas11}]], assparas11]
```
vereinfache reduzi··· ebne ein

```
True

True

True
```

The possible sign vectors are thus: {+,+,+,-} or {+,+,-,-}.
This implies exactly one zero in the positive interval for z and at most one admissible $\bar{s}$.

#### pole method:

##### Definition of the relevant expressions:

Poles are of the form $\bar{s}$=-r+s[i,j]. Their order is determined by the fitness order. Since the terms in the denominator with s[1,1] and s[1,3] are squared the sign does not change at the respective pole. For the others the sign changes.

```
pole11 = Limit[detStc, s̄ → -r + s[1, 1], Assumptions → r > 0];
```
Grenzwert Annahmen

```
pole13 = Limit[detStc, s̄ → -r + s[1, 3], Assumptions → r > 0];
```
Grenzwert Annahmen

```
pole12l = Limit[detStc, s̄ → -r + s[1, 2], Direction → 1, Assumptions → r > 0];
```
Grenzwert Richtung Annahmen

```
pole12r = Limit[detStc, s̄ → -r + s[1, 2], Direction → -1, Assumptions → r > 0];
```
Grenzwert Richtung Annahmen

```
pole21l = Limit[detStc, s̄ → -r + s[2, 1], Direction → 1, Assumptions → r > 0];
```
Grenzwert Richtung Annahmen

```
pole21r = Limit[detStc, s̄ → -r + s[2, 1], Direction → -1, Assumptions → r > 0];
```
Grenzwert Richtung Annahmen

```
pole22l = Limit[detStc, s̄ → -r, Direction → 1, Assumptions → r > 0];
```
Grenzwert Richtung Annahmen

```
pole22r = Limit[detStc, s̄ → -r, Direction → -1, Assumptions → r > 0];
```
Grenzwert Richtung Annahmen

```
sbar0 = FullSimplify[detStc /. s̄ → 0]
```
vereinfache vollständig

$$\frac{2\, r\, s[1,2]\, (-s[1,1] + s[1,3])\, s[2,1]}{(r - s[1,1])\, (r - s[1,2])\, (r - s[1,3])\, (r - s[2,1])}$$

Direction "-1" is the limit from right and "+1" from left.

In the limit $\bar{s}$->∞, the denominator is positive and thus the sign of the coefficient of $\bar{s}$^3 determines the sign of detStc.
In the limit $\bar{s}$->-∞, the denominator is negative and $\bar{s}$^3 is also negative, which implies that the sign of the coefficient of $\bar{s}$^3 determines the sign of detStc.

```
sbarinf = FullSimplify[Coefficient[Numerator[Together[detStc]], s̄, 3] / r^2];
```
vereinfache voll··· Koeffizient Zähler zusammen

## s22<s13<s21<s12<s11

$0 < r + \bar{s} - s[1, 1] < r + s[2, 1] - s[1, 1]$, since $\bar{s}$<s[2,1].

```
s22s13s21s12s11 =
 Simplify[Sign[{{sbarinf, pole22l}, {pole22r, pole13}, {pole13, pole21l},
 vereinfache Vorzeichen
    {pole21r, pole12l}, {pole12r, pole11}, {pole11, sbarinf}}],
  Flatten[{0 < s[1, 3] < s[2, 1] < s[1, 2] < s[1, 1], r > s[1, 1] - s[2, 1]}]]
  ebne ein
```

```
{{-Sign[s[1, 1] - 2 s[1, 2] + s[1, 3] - 2 s[2, 1]], 1}, {-1, -Sign[r - s[1, 3]]},
 {-Sign[r - s[1, 3]], 1}, {-1, -1}, {1, Sign[r - s[1, 1]]},
 {Sign[r - s[1, 1]], -Sign[s[1, 1] - 2 s[1, 2] + s[1, 3] - 2 s[2, 1]]}}
```

```
Simplify[s22s13s21s12s11, r > s[1, 1] > s[1, 3]]
vereinfache
Simplify[s22s13s21s12s11, s[1, 1] > r > s[1, 3]]
vereinfache
Simplify[s22s13s21s12s11, s[1, 1] > s[1, 3] > r]
vereinfache
```

```
{{-Sign[s[1, 1] - 2 s[1, 2] + s[1, 3] - 2 s[2, 1]], 1}, {-1, -1}, {-1, 1},
 {-1, -1}, {1, 1}, {1, -Sign[s[1, 1] - 2 s[1, 2] + s[1, 3] - 2 s[2, 1]]}}
```

```
{{-Sign[s[1, 1] - 2 s[1, 2] + s[1, 3] - 2 s[2, 1]], 1}, {-1, -1}, {-1, 1},
 {-1, -1}, {1, -1}, {-1, -Sign[s[1, 1] - 2 s[1, 2] + s[1, 3] - 2 s[2, 1]]}}
```

```
{{-Sign[s[1, 1] - 2 s[1, 2] + s[1, 3] - 2 s[2, 1]], 1}, {-1, 1}, {1, 1},
 {-1, -1}, {1, -1}, {-1, -Sign[s[1, 1] - 2 s[1, 2] + s[1, 3] - 2 s[2, 1]]}}
```

If r>s[1,1], then sign distribution is not clear, however all poles are negative, therefore:

```
s22s13s21s12s110incl =
 Simplify[Sign[{{sbarinf, pole22l}, {pole22r, pole13}, {pole13, pole21l},
 vereinfache Vorzeichen
    {pole21r, pole12l}, {pole12r, pole11}, {pole11, sbar0}, {sbar0, sbarinf}}],
  Flatten[{0 < s[1, 3] < s[1, 1] < r, 0 < s[1, 3] < s[2, 1] < s[1, 2] < s[1, 1],
  ebne ein
    r > s[1, 1] - s[2, 1]}]]
Simplify[Sign[{{sbarinf, pole22l}, {pole22r, pole13}, {pole13, pole21l},
vereinfache Vorzeichen
   {pole21r, pole12l}, {pole12r, pole11}, {pole11, sbar0}, {sbar0, sbarinf}}],
 Flatten[{0 < r < s[1, 3] < s[1, 1], 0 < s[1, 3] < s[2, 1] < s[1, 2] < s[1, 1],
 ebne ein
   r > s[1, 1] - s[2, 1]}]]
```

```
{{-Sign[s[1, 1] - 2 s[1, 2] + s[1, 3] - 2 s[2, 1]], 1}, {-1, -1}, {-1, 1},
 {-1, -1}, {1, 1}, {1, -1}, {-1, -Sign[s[1, 1] - 2 s[1, 2] + s[1, 3] - 2 s[2, 1]]}}
```

```
{{1, 1}, {-1, 1}, {1, 1}, {-1, -1}, {1, -1}, {-1, -1}, {-1, 1}}
```

In the relevant interval (pole11,sbar0) is at most one zero.

## s13<s22<s21<s12<s11

$0 < r + \bar{s} - s[1, 1] < r + s[2, 1] - s[1, 1]$, since $\bar{s}$<s[2,1].

```
s13s22s21s12s11 =
 Simplify[Sign[{{sbarinf, pole13}, {pole13, pole22l}, {pole22r, pole21l},
 vereinfache Vorzeichen
    {pole21r, pole12l}, {pole12r, pole11}, {pole11, sbarinf}}],
  Flatten[{s[1, 3] < 0 < s[2, 1] < s[1, 2] < s[1, 1], r > s[1, 1] - s[2, 1]}]]
  ebne ein
```

```
{{-Sign[s[1, 1] - 2 s[1, 2] + s[1, 3] - 2 s[2, 1]], 1},
 {1, Sign[2 s[1, 1] s[1, 3] + r (s[1, 1] + s[1, 3])]},
 {-Sign[2 s[1, 1] s[1, 3] + r (s[1, 1] + s[1, 3])], 1}, {-1, -1}, {1, Sign[r - s[1, 1]]},
 {Sign[r - s[1, 1]], -Sign[s[1, 1] - 2 s[1, 2] + s[1, 3] - 2 s[2, 1]]}}
```

```
Simplify[s13s22s21s12s11, r < s[1, 1]]
vereinfache
```

```
{{-Sign[s[1, 1] - 2 s[1, 2] + s[1, 3] - 2 s[2, 1]], 1},
 {1, Sign[2 s[1, 1] s[1, 3] + r (s[1, 1] + s[1, 3])]},
 {-Sign[2 s[1, 1] s[1, 3] + r (s[1, 1] + s[1, 3])], 1}, {-1, -1},
 {1, -1}, {-1, -Sign[s[1, 1] - 2 s[1, 2] + s[1, 3] - 2 s[2, 1]]}}
```

```
s13s22s21s12s11incl0 =
 Simplify[Sign[{{sbarinf, pole13}, {pole13, pole22l}, {pole22r, pole21l},
 vereinfache Vorzeichen
    {pole21r, pole12l}, {pole12r, pole11}, {pole11, sbar0}, {sbar0, sbarinf}}],
  Flatten[{s[1, 3] < 0 < s[1, 1] < r, s[1, 3] < 0 < s[2, 1] < s[1, 2] < s[1, 1],
  ebne ein
    r > s[1, 1] - s[2, 1]}]]
```

```
{{-Sign[s[1, 1] - 2 s[1, 2] + s[1, 3] - 2 s[2, 1]], 1},
 {1, Sign[2 s[1, 1] s[1, 3] + r (s[1, 1] + s[1, 3])]},
 {-Sign[2 s[1, 1] s[1, 3] + r (s[1, 1] + s[1, 3])], 1}, {-1, -1},
 {1, 1}, {1, -1}, {-1, -Sign[s[1, 1] - 2 s[1, 2] + s[1, 3] - 2 s[2, 1]]}}
```

There are three intervals with a sign change regardless of the order of r and s[1,1].

## s22<s13<s11<s21<s12

$0 < r + \bar{s} - s[1, 2] < r+s[2,1]-s[1,2]$

```
s22s13s11s21s12 =
 Simplify[Sign[{{sbarinf, pole22l}, {pole22r, pole13}, {pole13, pole11},
 vereinfache Vorzeichen
    {pole11, pole21l}, {pole21r, pole12l}, {pole12r, sbarinf}}],
  Flatten[{0 < s[1, 3] < s[1, 1] < s[2, 1] < s[1, 2], r > s[1, 2] - s[2, 1]}]]
  ebne ein
```

```
{{1, 1}, {-1, -Sign[r - s[1, 3]]},
 {-Sign[r - s[1, 3]], Sign[r - s[1, 1]]}, {Sign[r - s[1, 1]], -1}, {1, 1}, {-1, 1}}
```

```
Simplify[s22s13s11s21s12, r > s[1, 1]]
```
vereinfache
```
Simplify[s22s13s11s21s12, s[1, 3] < r < s[1, 1]]
```
vereinfache
```
Simplify[s22s13s11s21s12, r < s[1, 3] < s[1, 1]]
```
vereinfache

```
{{1, 1}, {-1, -Sign[r - s[1, 3]]}, {-Sign[r - s[1, 3]], 1}, {1, -1}, {1, 1}, {-1, 1}}

{{1, 1}, {-1, -1}, {-1, -1}, {-1, -1}, {1, 1}, {-1, 1}}

{{1, 1}, {-1, 1}, {1, -1}, {-1, -1}, {1, 1}, {-1, 1}}
```

Not clear for s[1,3]<r<s[1,1], which implies $-r + s[1, 3] < 0$<-r+s[1,1]:

```
s22s13s11s21s12incl0 =
 Simplify[Sign[{{sbarinf, pole22l}, {pole22r, pole13}, {pole13, sbar0},
    {sbar0, pole11}, {pole11, pole21l}, {pole21r, pole12l}, {pole12r, sbarinf}}],
  Flatten[{0 < s[1, 3] < r < s[1, 1] < s[2, 1] < s[1, 2], r > s[1, 2] - s[2, 1]}]]
```
vereinfache Vorzeichen ebne ein

```
{{1, 1}, {-1, -1}, {-1, 1}, {1, -1}, {-1, -1}, {1, 1}, {-1, 1}}
```

two more zeroes are found and the claim is proven.

## s22<s13<s21<s11<s12

$0 < r + \bar{s} - s[1, 2] < r + s[2, 1] - s[1, 2]$

```
s22s13s21s11s12 =
 Simplify[Sign[{{sbarinf, pole22l}, {pole22r, pole13}, {pole13, pole21l},
    {pole21r, pole11}, {pole11, pole12l}, {pole12r, sbarinf}}],
  Flatten[{0 < s[1, 3] < s[2, 1] < s[1, 1] < s[1, 2], r > s[1, 2] - s[2, 1]}]]
```
vereinfache Vorzeichen ebne ein

```
{{1, 1}, {-1, -Sign[r - s[1, 3]]}, {-Sign[r - s[1, 3]], 1},
 {-1, Sign[r - s[1, 1]]}, {Sign[r - s[1, 1]], 1}, {-1, 1}}
```

There are three distinct intervals that contain each one zero.

## s13<s22<s11<s21<s12

$0 < r + \bar{s} - s[1, 2] < r + s[2, 1] - s[1, 2]$

```
s13s22s11s21s12 =
 Simplify[Sign[{{sbarinf, pole13}, {pole13, pole22l}, {pole22r, pole11},
    {pole11, pole21l}, {pole21r, pole12l}, {pole12r, sbarinf}}],
  Flatten[{s[1, 3] < 0 < s[1, 1] < s[2, 1] < s[1, 2], r > s[1, 2] - s[2, 1]}]]
```
vereinfache Vorzeichen ebne ein

```
{{1, 1}, {1, Sign[2 s[1, 1] s[1, 3] + r (s[1, 1] + s[1, 3])]},
 {-Sign[2 s[1, 1] s[1, 3] + r (s[1, 1] + s[1, 3])], Sign[r - s[1, 1]]},
 {Sign[r - s[1, 1]], -1}, {1, 1}, {-1, 1}}
```

```
Simplify[s13s22s11s21s12, s[1, 1] < r]
vereinfache
Simplify[s13s22s11s21s12, r < s[1, 1]]
vereinfache
```

```
{{1, 1}, {1, Sign[2 s[1, 1] s[1, 3] + r (s[1, 1] + s[1, 3])]},
 {-Sign[2 s[1, 1] s[1, 3] + r (s[1, 1] + s[1, 3])], 1}, {1, -1}, {1, 1}, {-1, 1}}
```

```
{{1, 1}, {1, Sign[2 s[1, 1] s[1, 3] + r (s[1, 1] + s[1, 3])]},
 {-Sign[2 s[1, 1] s[1, 3] + r (s[1, 1] + s[1, 3])], -1}, {-1, -1}, {1, 1}, {-1, 1}}
```

It is not clear yet for 0<r<s[1,1], however we can look at the sign at sbar=0:

```
s13s22s11s21s12incl0 =
 Simplify[Sign[{{sbarinf, pole13}, {pole13, pole22l}, {pole22r, sbar0},
 vereinfache Vorzeichen
    {sbar0, pole11}, {pole11, pole21l}, {pole21r, pole12l}, {pole12r, sbarinf}}],
  Flatten[{s[1, 3] < 0 < r < s[1, 1] < s[2, 1] < s[1, 2], r > s[1, 2] - s[2, 1]}]]
  ebne ein
```

```
{{1, 1}, {1, Sign[2 s[1, 1] s[1, 3] + r (s[1, 1] + s[1, 3])]},
 {-Sign[2 s[1, 1] s[1, 3] + r (s[1, 1] + s[1, 3])], 1}, {1, -1}, {-1, -1}, {1, 1}, {-1, 1}}
```

Each case has three distinct intervals which contain a zero.

## s13<s22<s21<s11<s12

$0 < r + \overline{s} - s[1, 2] < r + s[2, 1] - s[1, 2]$

```
s13s22s21s11s12 =
 Simplify[Sign[{{sbarinf, pole13}, {pole13, pole22l}, {pole22r, pole21l},
 vereinfache Vorzeichen
    {pole21r, pole11}, {pole11, pole12l}, {pole12r, sbarinf}}],
  Flatten[{s[1, 3] < 0 < s[2, 1] < s[1, 1] < s[1, 2], r > s[1, 2] - s[2, 1]}]]
  ebne ein
```

```
{{1, 1}, {1, Sign[2 s[1, 1] s[1, 3] + r (s[1, 1] + s[1, 3])]},
 {-Sign[2 s[1, 1] s[1, 3] + r (s[1, 1] + s[1, 3])], 1},
 {-1, Sign[r - s[1, 1]]}, {Sign[r - s[1, 1]], 1}, {-1, 1}}
```

There are three distinct intervals where each contains a zero.

## s13<s11<s22<s21<s12

$0 < r + \overline{s} - s[1, 2] < r + s[2, 1] - s[1, 2]$

```
s13s11s22s21s12 = Simplify[Sign[{{sbarinf, pole13}, {pole13, pole11},
                  vereinfache Vorzeichen
    {pole11, pole22l}, {pole22r, pole21l}, {pole21r, pole12l}, {pole12r, sbarinf}}],
  Flatten[{s[1, 3] < s[1, 1] < 0 < s[2, 1] < s[1, 2], r > s[1, 2] - s[2, 1]}]]
  ebne ein
```

```
{{1, 1}, {1, -1}, {-1, Sign[2 s[1, 1] s[1, 3] + r (s[1, 1] + s[1, 3])]},
 {-Sign[2 s[1, 1] s[1, 3] + r (s[1, 1] + s[1, 3])], -1}, {1, 1}, {-1, 1}}
```

There are three distinct intervals where each contains a zero.

### Internal equilibrium coordinates:

We scale the fitness matrix such that $\bar{s} = 0$.

```
S̃c = FullSimplify[S̃[3] /. centrosym[3] /. s̄ → 0];
```
vereinfache vollständig

```
MatrixForm[S̃c]
```
Matritzenform

$$\begin{pmatrix} \frac{s[1,1]}{r-s[1,1]} & \frac{s[1,2]}{r-s[1,2]} & \frac{s[1,3]}{r-s[1,3]} \\ \frac{s[2,1]}{r-s[2,1]} & \frac{s[2,2]}{r-s[2,2]} & \frac{s[2,1]}{r-s[2,1]} \\ \frac{s[1,3]}{r-s[1,3]} & \frac{s[1,2]}{r-s[1,2]} & \frac{s[1,1]}{r-s[1,1]} \end{pmatrix}$$

```
detStildec = FullSimplify[Det[S̃c]]
```
vereinfache voll··· Determinant

```
(r^2 (-s[1, 1] + s[1, 3]) (2 (r - s[1, 1]) s[1, 2] (r - s[1, 3]) s[2, 1] +
    (-(r - s[1, 2]) (r s[1, 1] + (r - 2 s[1, 1]) s[1, 3]) + (r s[1, 1] - 2 r s[1, 2] +
        s[1, 1] s[1, 2] + (r - 2 s[1, 1] + s[1, 2]) s[1, 3]) s[2, 1]) s[2, 2])) /
 ((r - s[1, 1])^2 (r - s[1, 2]) (r - s[1, 3])^2 (r - s[2, 1]) (r - s[2, 2]))
```

We solve the Determinant of $\tilde{S}_c$ for s[2,2], since it is linear in s[2,2]:

```
š = FullSimplify[s[2, 2] /. Solve[detStildec == 0, s[2, 2]]][[1]]
```
vereinfache vollständig löse

```
(2 (r - s[1, 1]) s[1, 2] (r - s[1, 3]) s[2, 1]) /
 ((r - s[1, 2]) (r s[1, 1] + (r - 2 s[1, 1]) s[1, 3]) -
   (r s[1, 1] - 2 r s[1, 2] + s[1, 1] s[1, 2] + (r - 2 s[1, 1] + s[1, 2]) s[1, 3]) s[2, 1])
```

```
š = (2 (r - s[1, 1]) s[1, 2] (r - s[1, 3]) s[2, 1]) /
    ((r - s[1, 2]) (r s[1, 1] + (r - 2 s[1, 1]) s[1, 3]) -
      (r s[1, 1] - 2 r s[1, 2] + s[1, 1] s[1, 2] + (r - 2 s[1, 1] + s[1, 2]) s[1, 3]) s[2, 1]);
```

If we define F and E1 (E is a built-in function in Mathematica) as in the main text, the expression for š holds.

```
FE1tr = {F → s[1, 1] + s[1, 3] - 2 r, E1 → s[1, 1] (r - s[1, 3]) + s[1, 3] (r - s[1, 1])};
```

```
FullSimplify[š == (2 s[1, 2] s[2, 1] (r - s[1, 1]) (r - s[1, 3])) /
```
vereinfache vollständig
```
    ((r - s[1, 2]) E1 - s[2, 1] (E1 + s[1, 2] F)) /. FE1tr]
```

```
True
```

```
Stildec = FullSimplify[S̃c /. s[2, 2] → š];
```
vereinfache vollständig

```
Det[Stildec]
```
Determinante

```
0
```

This can be solved to get the mean allele freqencies at the equilibrium. (Equ. 16b and c from the main text)

```
ptr0 =
  FullSimplify[Solve[{p[1], p[2], 1 - p[1] - p[2]}.Stildec == 0, {p[1], p[2]}][[1]]];
  vereinfache voll··· löse
ptr = Join[ptr0, {p[3] → FullSimplify[1 - p[1] - p[2] /. ptr0]}]
      verknüpfe                vereinfache vollständig
```

```
{p[1] → -(((r - s[1, 1]) (r - s[1, 3]) s[2, 1]) /
      (r (r s[1, 1] + r s[1, 3] - 2 s[1, 1] s[1, 3] + (-2 r + s[1, 1] + s[1, 3]) s[2, 1]))),
 p[2] → ((r s[1, 1] + (r - 2 s[1, 1]) s[1, 3]) (r - s[2, 1])) /
    (r (r s[1, 1] + r s[1, 3] - 2 s[1, 1] s[1, 3] + (-2 r + s[1, 1] + s[1, 3]) s[2, 1])),
 p[3] → -(((r - s[1, 1]) (r - s[1, 3]) s[2, 1]) /
      (r (r s[1, 1] + r s[1, 3] - 2 s[1, 1] s[1, 3] + (-2 r + s[1, 1] + s[1, 3]) s[2, 1])))}
```

```
qtr0 =
  FullSimplify[Solve[Stildec.{q[1], q[2], 1 - q[1] - q[2]} == 0, {q[1], q[2]}][[1]]];
  vereinfache voll··· löse
qtr = Join[qtr0, {q[3] → FullSimplify[1 - q[1] - q[2] /. qtr0]}]
      verknüpfe                vereinfache vollständig
```

```
{q[1] → -(((r - s[1, 1]) s[1, 2] (r - s[1, 3])) /
      (r (r s[1, 1] - 2 r s[1, 2] + s[1, 1] s[1, 2] + (r - 2 s[1, 1] + s[1, 2]) s[1, 3]))),
 q[2] → ((r - s[1, 2]) (r s[1, 1] + (r - 2 s[1, 1]) s[1, 3])) /
    (r (r s[1, 1] - 2 r s[1, 2] + s[1, 1] s[1, 2] + (r - 2 s[1, 1] + s[1, 2]) s[1, 3])),
 q[3] → -(((r - s[1, 1]) s[1, 2] (r - s[1, 3])) /
      (r (r s[1, 1] - 2 r s[1, 2] + s[1, 1] s[1, 2] + (r - 2 s[1, 1] + s[1, 2]) s[1, 3])))}
```

Also the denominators of the expressions q[j] are the same for all j and also for p[i]. The numerators are simpler to check but also coincide with the expressions given in eq. (37) of the main text.

```
FullSimplify[Denominator[q[3] /. qtr] == r (E1 + s[1, 2] F) /. FE1tr]
vereinfache voll··· Nenner
```

```
True
```

Using formula for the equilibrium coordinates eq. (17), we compute the coordinates for the equilibrium with mean fitness sclaed to zero:

```
pijequi = Flatten[Table[freqarray[3, 3][[i, j]] →
          ebne ein   Tabelle
    FullSimplify[r {p[1], p[2], p[3]}[[i]] {q[1], q[2], q[3]}[[j]] / (r - s[i, j]) /.
    vereinfache vollständig
        centrosym[3] /. s[2, 2] → š /. qtr /. ptr], {i, 3}, {j, 3}]]
{p[1, 1] → ((r - s[1, 1]) s[1, 2] (r - s[1, 3])^2 s[2, 1]) /
   (r (r s[1, 1] - 2 r s[1, 2] + s[1, 1] s[1, 2] + (r - 2 s[1, 1] + s[1, 2]) s[1, 3])
     (r s[1, 1] + r s[1, 3] - 2 s[1, 1] s[1, 3] + (-2 r + s[1, 1] + s[1, 3]) s[2, 1])),
 p[1, 2] → -(((r - s[1, 1]) (r - s[1, 3]) (r s[1, 1] + (r - 2 s[1, 1]) s[1, 3]) s[2, 1]) /
     (r (r s[1, 1] - 2 r s[1, 2] + s[1, 1] s[1, 2] + (r - 2 s[1, 1] + s[1, 2]) s[1, 3])
       (r s[1, 1] + r s[1, 3] - 2 s[1, 1] s[1, 3] + (-2 r + s[1, 1] + s[1, 3]) s[2, 1]))),
 p[1, 3] → ((r - s[1, 1])^2 s[1, 2] (r - s[1, 3]) s[2, 1]) /
   (r (r s[1, 1] - 2 r s[1, 2] + s[1, 1] s[1, 2] + (r - 2 s[1, 1] + s[1, 2]) s[1, 3])
     (r s[1, 1] + r s[1, 3] - 2 s[1, 1] s[1, 3] + (-2 r + s[1, 1] + s[1, 3]) s[2, 1])),
 p[2, 1] → -(((r - s[1, 1]) s[1, 2] (r - s[1, 3]) (r s[1, 1] + (r - 2 s[1, 1]) s[1, 3])) /
     (r (r s[1, 1] - 2 r s[1, 2] + s[1, 1] s[1, 2] + (r - 2 s[1, 1] + s[1, 2]) s[1, 3])
       (r s[1, 1] + r s[1, 3] - 2 s[1, 1] s[1, 3] + (-2 r + s[1, 1] + s[1, 3]) s[2, 1]))),
 p[2, 2] → ((r s[1, 1] + (r - 2 s[1, 1]) s[1, 3])
     ((r - s[1, 2]) (r s[1, 1] + (r - 2 s[1, 1]) s[1, 3]) - (r s[1, 1] - 2 r s[1, 2] +
          s[1, 1] s[1, 2] + (r - 2 s[1, 1] + s[1, 2]) s[1, 3]) s[2, 1])) /
   (r (r s[1, 1] - 2 r s[1, 2] + s[1, 1] s[1, 2] + (r - 2 s[1, 1] + s[1, 2]) s[1, 3])
     (r s[1, 1] + r s[1, 3] - 2 s[1, 1] s[1, 3] + (-2 r + s[1, 1] + s[1, 3]) s[2, 1])),
 p[2, 3] → -(((r - s[1, 1]) s[1, 2] (r - s[1, 3]) (r s[1, 1] + (r - 2 s[1, 1]) s[1, 3])) /
     (r (r s[1, 1] - 2 r s[1, 2] + s[1, 1] s[1, 2] + (r - 2 s[1, 1] + s[1, 2]) s[1, 3])
       (r s[1, 1] + r s[1, 3] - 2 s[1, 1] s[1, 3] + (-2 r + s[1, 1] + s[1, 3]) s[2, 1]))),
 p[3, 1] → ((r - s[1, 1])^2 s[1, 2] (r - s[1, 3]) s[2, 1]) /
   (r (r s[1, 1] - 2 r s[1, 2] + s[1, 1] s[1, 2] + (r - 2 s[1, 1] + s[1, 2]) s[1, 3])
     (r s[1, 1] + r s[1, 3] - 2 s[1, 1] s[1, 3] + (-2 r + s[1, 1] + s[1, 3]) s[2, 1])),
 p[3, 2] → -(((r - s[1, 1]) (r - s[1, 3]) (r s[1, 1] + (r - 2 s[1, 1]) s[1, 3]) s[2, 1]) /
     (r (r s[1, 1] - 2 r s[1, 2] + s[1, 1] s[1, 2] + (r - 2 s[1, 1] + s[1, 2]) s[1, 3])
       (r s[1, 1] + r s[1, 3] - 2 s[1, 1] s[1, 3] + (-2 r + s[1, 1] + s[1, 3]) s[2, 1]))),
 p[3, 3] → ((r - s[1, 1]) s[1, 2] (r - s[1, 3])^2 s[2, 1]) /
   (r (r s[1, 1] - 2 r s[1, 2] + s[1, 1] s[1, 2] + (r - 2 s[1, 1] + s[1, 2]) s[1, 3])
     (r s[1, 1] + r s[1, 3] - 2 s[1, 1] s[1, 3] + (-2 r + s[1, 1] + s[1, 3]) s[2, 1]))}
```

### Explicit expressions in the new coordinates:

Representation of u in p-coordinates.

```
uretra = {u1 -> freqlist[3, 3][[1]] - freqlist[3, 3][[9]],
   u2 -> freqlist[3, 3][[2]] - freqlist[3, 3][[8]],
   u3 -> freqlist[3, 3][[3]] - freqlist[3, 3][[7]],
   u4 -> freqlist[3, 3][[4]] - freqlist[3, 3][[6]],
   u5 -> freqlist[3, 3][[1]] +
     freqlist[3, 3][[9]] - freqlist[3, 3][[2]] - freqlist[3, 3][[8]],
   u6 -> freqlist[3, 3][[3]] + freqlist[3, 3][[7]] -
     freqlist[3, 3][[4]] - freqlist[3, 3][[6]],
   u7 -> freqlist[3, 3][[1]] + freqlist[3, 3][[9]] + freqlist[3, 3][[2]] +
     freqlist[3, 3][[8]] - freqlist[3, 3][[3]] - freqlist[3, 3][[7]] -
     freqlist[3, 3][[4]] - freqlist[3, 3][[6]],
   u8 -> freqlist[3, 3][[5]]};
```

```
ucoor = {u1, u2, u3, u4, u5, u6, u7, u8};
```

Representation of p in u-coordinates.

$$\text{utra} = \{p[1,1] \to \frac{1}{8}(1 + 4\,u1 + 2\,u5 + u7 - u8),$$
$$p[1,2] \to \frac{1}{8}(1 + 4\,u2 - 2\,u5 + u7 - u8),\ p[1,3] \to \frac{1}{8}(1 + 4\,u3 + 2\,u6 - u7 - u8),$$
$$p[2,1] \to \frac{1}{8}(1 + 4\,u4 - 2\,u6 - u7 - u8),\ p[2,2] \to u8,$$
$$p[2,3] \to \frac{1}{8}(1 - 4\,u4 - 2\,u6 - u7 - u8),\ p[3,1] \to \frac{1}{8}(1 - 4\,u3 + 2\,u6 - u7 - u8),$$
$$p[3,2] \to \frac{1}{8}(1 - 4\,u2 - 2\,u5 + u7 - u8),\ p[3,3] \to \frac{1}{8}(1 - 4\,u1 + 2\,u5 + u7 - u8)\};$$

mean fitness and simplex condition in u coordinates:

```
sbaru = Collect[sbar[3, 3] /. utra /. centrosym[3], fitlist[3, 3], FullSimplify]
         gruppiere Koeffizienten                                vereinfache vollstär
FullSimplify[simplex[3, 3] /. utra]
vereinfache vollständig
```

$$\frac{1}{4}(1 + 2\,u5 + u7 - u8)\,s[1,1] + \frac{1}{4}(1 - 2\,u5 + u7 - u8)\,s[1,2] +$$
$$\frac{1}{4}(1 + 2\,u6 - u7 - u8)\,s[1,3] + \frac{1}{4}(1 - 2\,u6 - u7 - u8)\,s[2,1] + u8\,s[2,2]$$

{True}

Then we determine the ODE in u-coordinates:

```
ulistf = FullSimplify[
         vereinfache vollständig
   ucoor /. uretra /. Table[freqlist[3, 3][[i]] → listsys[3, 3][[i]], {i, 9}] /.
                      Tabelle
      centrosym[3] /. utra /. sbarformal → sbaruformal];
ulist = FullSimplify[ulistf /. sbaruformal → sbaru]
        vereinfache vollständig
```

```
{1/8 (r (2 u3 (u5 - u6 - u7) +
        2 u1 (-1 + u5 + u6 - 3 u8) + u2 (3 + 2 u5 - u7 - 3 u8) + u4 (3 + 2 u6 + u7 - 3 u8)) -
     2 u1 ((-3 + 2 u5 + u7 - u8) s[1, 1] + (1 - 2 u5 + u7 - u8) s[1, 2] + (1 + 2 u6) s[1, 3] +
        (1 - 2 u6) s[2, 1] - (u7 + u8) (s[1, 3] + s[2, 1]) + 4 u8 s[2, 2])),
 1/4 (r (-3 u2 + u3 - (u2 + u3) (2 u5 - u7 - 3 u8) + u1 (1 - 2 u5 + u7 + 3 u8)) +
     u2 (-(1 + 2 u5 + u7 - u8) s[1, 1] + (3 + 2 u5 - u7 + u8) s[1, 2] - (1 + 2 u6) s[1, 3] +
        (-1 + 2 u6) s[2, 1] + (u7 + u8) (s[1, 3] + s[2, 1]) - 4 u8 s[2, 2])),
 1/8 (r (2 u1 (u5 - u6 - u7) + 2 u3 (-1 + u5 + u6 - 3 u8) + u2 (3 + 2 u5 - u7 - 3 u8) -
        u4 (3 + 2 u6 + u7 - 3 u8)) - 2 u3 ((1 + 2 u5 + u7 - u8) s[1, 1] + (1 - 2 u5 + u7 - u8) s[1, 2] +
        (-3 + 2 u6) s[1, 3] + (1 - 2 u6) s[2, 1] - (u7 + u8) (s[1, 3] + s[2, 1]) + 4 u8 s[2, 2])),
 1/4 (-r (u3 + 3 u4 + u1 (-1 + 2 u6 + u7) - (u3 - u4) (2 u6 + u7)) + 3 r (u1 - u3 + u4) u8 +
     u4 (-(1 + 2 u5 + u7 - u8) s[1, 1] + (-1 + 2 u5 - u7 + u8) s[1, 2] - (1 + 2 u6) s[1, 3] +
        (3 + 2 u6) s[2, 1] + (u7 + u8) (s[1, 3] + s[2, 1]) - 4 u8 s[2, 2])),
 1/32 (r (16 (u1 + u2 + u3) (u1 - u3 + u4) + 2 u5 (-7 + 6 u6 + 3 u7 - 9 u8) -
        (3 + 2 u6 + u7 - 3 u8) (-1 + 3 u7 + 9 u8)) +
     8 (-(-1 + u5) (1 + 2 u5 + u7 - u8) s[1, 1] + (1 + u5) (-1 + 2 u5 - u7 + u8) s[1, 2] +
        u5 ((-1 - 2 u6 + u7 + u8) s[1, 3] + (-1 + 2 u6 + u7 + u8) s[2, 1] - 4 u8 s[2, 2]))),
 1/32 (-r (16 (u1 + u2 + u3) (u1 - u3 + u4) - 2 u5 (1 + 6 u6 + 3 u7 - 9 u8) +
        (1 + 3 u7 - 9 u8) (-3 + u7 + 3 u8) + 2 u6 (7 + 3 u7 + 9 u8)) -
     8 (2 u6^2 (s[1, 3] - s[2, 1]) + (-1 + u7 + u8) (s[1, 3] - s[2, 1]) +
        u6 ((1 + 2 u5 + u7 - u8) s[1, 1] + (1 - 2 u5 + u7 - u8) s[1, 2] -
           (1 + u7 + u8) (s[1, 3] + s[2, 1]) + 4 u8 s[2, 2]))),
 1/4 (2 r (2 (u1 + u2 + u3) (u1 - u3 + u4) - u5 + u6 - u7) - (-1 + u7) (1 + 2 u5 + u7 - u8) s[1, 1] +
     s[1, 2] - 2 u5 s[1, 2] + 2 u5 u7 s[1, 2] - u7^2 s[1, 2] - u8 s[1, 2] +
     u7 u8 s[1, 2] - (1 + 2 u6) s[1, 3] - 2 u6 u7 s[1, 3] + u7^2 s[1, 3] +
     u8 s[1, 3] + u7 u8 s[1, 3] - s[2, 1] + 2 u6 s[2, 1] + 2 u6 u7 s[2, 1] +
     u7^2 s[2, 1] + u8 s[2, 1] + u7 u8 s[2, 1] - 4 u7 u8 s[2, 2]),
 1/16 (r ((-1 + 2 u5 - u7) (-1 + 2 u6 + u7) - 2 (5 + 3 u5 + 3 u6) u8 + 9 u8^2) +
     4 u8 (-(1 + 2 u5 + u7 - u8) s[1, 1] + (-1 + 2 u5 - u7 + u8) s[1, 2] +
        (-1 - 2 u6 + u7 + u8) s[1, 3] + (-1 + 2 u6 + u7 + u8) s[2, 1] + 4 s[2, 2] - 4 u8 s[2, 2]))}
```

```
ulist={1/8 (r (2 u3 (u5-u6-u7)+2 u1 (-1+u5+u6-3 u8)+u2 (3+2 u5-u7-3 u8)+u4 (3+2 u6+u7-3 u8))-
```

and similarly the equilibrium in u-coordinates:

```
ueq = FullSimplify[uretra /. pijequi]
        vereinfache vollständig
```

```
{u1 → p[1, 1] - p[3, 3], u2 → p[1, 2] - p[3, 2], u3 → p[1, 3] - p[3, 1], u4 → p[2, 1] - p[2, 3],
  u5 → p[1, 1] - p[1, 2] - p[3, 2] + p[3, 3], u6 → p[1, 3] - p[2, 1] - p[2, 3] + p[3, 1],
  u7 → p[1, 1] + p[1, 2] - p[1, 3] - p[2, 1] - p[2, 3] - p[3, 1] + p[3, 2] + p[3, 3],
  u8 → p[2, 2]} /. pijequi
```

it is indeed a solution:

```
Simplify[ulist /. ueq /. s[2, 2] → š]
vereinfache
```

```
{0, 0, 0, 0, 0, 0, 0, 0}
```

Jacobian at u1=u2=u3=u4=0:

```
Ju0 = Simplify[D[ulist, {ucoor}]];
      vereinfache leite ab
MatrixForm[FullSimplify[Ju0 /. {u1 → 0, u2 → 0, u3 → 0, u4 → 0}]]
Matritzenform vereinfache vollständig
```

```
( 1/8 (2 r (-1 + u5 + u6 - 3 u8) - 2 ((-3 + 2 u5 + u7 - u8) s[1, 1] + (1 - 2 u5 + u7 - u8) s[1, 2] + (1 + 2 u
                                                                       1/4 r (1 - 2 u5 + u7 + 3 u
                                                                         1/4 r (u5 - u6 - u7)
                                                                    - 1/4 r (-1 + 2 u6 + u7 - 3
                                                                                0
                                                                                0
                                                                                0
                                                                                0
```

sbaru is zero at equilibrium:

```
Ju1 = Ju0;
Table[Ju1[[i, i]] = FullSimplify[Ju0[[i, i]] + sbaru], {i, 8}]
Tabelle                    vereinfache vollständig
```

```
{1/4 r (-1 + u5 + u6 - 3 u8) + s[1, 1], 1/4 (r (-3 - 2 u5 + u7 + 3 u8) + 4 s[1, 2]),
 1/4 r (-1 + u5 + u6 - 3 u8) + s[1, 3], - 1/4 r (3 + 2 u6 + u7 - 3 u8) + s[2, 1],
 1/16 (r (-7 + 6 u6 + 3 u7 - 9 u8) - 8 (-1 + u5) s[1, 1] + 8 (1 + u5) s[1, 2]),
 1/16 (r (-7 + 6 u5 - 3 u7 - 9 u8) - 8 (-1 + u6) s[1, 3] + 8 (1 + u6) s[2, 1]),
 1/4 (-2 r - (-1 + u7) s[1, 1] - (-1 + u7) s[1, 2] + (1 + u7) (s[1, 3] + s[2, 1])),
 1/8 (-r (5 + 3 u5 + 3 u6 - 9 u8) +
    2 u8 (s[1, 1] + s[1, 2] + s[1, 3] + s[2, 1]) - 8 (-1 + u8) s[2, 2])}
```

```
Ju2 = FullSimplify[Ju1 /. {u1 → 0, u2 → 0, u3 → 0, u4 → 0}];
```
vereinfache vollständig

```
MatrixForm[Ju2]
```
Matritzenform

| | | |
|---|---|---|
| $\frac{1}{4}$ r (-1 + u5 + u6 - 3 u8) + s[1, 1] | $\frac{1}{8}$ r (3 + 2 u5 - u7 - 3 u8) | $\frac{1}{4}$ r (u5 - u6 - [illegible] |
| $\frac{1}{4}$ r (1 - 2 u5 + u7 + 3 u8) | $\frac{1}{4}$ (r (-3 - 2 u5 + u7 + 3 u8) + 4 s[1, 2]) | $\frac{1}{4}$ r (1 - 2 u5 + u7 [illegible] |
| $\frac{1}{4}$ r (u5 - u6 - u7) | $\frac{1}{8}$ r (3 + 2 u5 - u7 - 3 u8) | $\frac{1}{4}$ r (-1 + u5 + u6 - 3 u[illegible] |
| -$\frac{1}{4}$ r (-1 + 2 u6 + u7 - 3 u8) | 0 | $\frac{1}{4}$ r (-1 + 2 u6 + u7 [illegible] |
| 0 | 0 | 0 |
| 0 | 0 | 0 |
| 0 | 0 | 0 |
| 0 | 0 | 0 |

JuA is the first block $C_1$ from the main text:

```
JuA = FullSimplify[Take[Ju2, 4, 4]];
```
vereinfache voll··· entferne

```
JuA = {{1/4 r (-1 + u5 + u6 - 3 u8) + s[1, 1], 1/8 r (3 + 2 u5 - u7 - 3 u8),
    1/4 r (u5 - u6 - u7), 1/8 r (3 + 2 u6 + u7 - 3 u8)}, {1/4 r (1 - 2 u5 + u7 + 3 u8),
    1/4 (r (-3 - 2 u5 + u7 + 3 u8) + 4 s[1, 2]), 1/4 r (1 - 2 u5 + u7 + 3 u8), 0},
   {1/4 r (u5 - u6 - u7), 1/8 r (3 + 2 u5 - u7 - 3 u8), 1/4 r (-1 + u5 + u6 - 3 u8) + s[1, 3],
    -1/8 r (3 + 2 u6 + u7 - 3 u8)}, {-1/4 r (-1 + 2 u6 + u7 - 3 u8), 0,
    1/4 r (-1 + 2 u6 + u7 - 3 u8), -1/4 r (3 + 2 u6 + u7 - 3 u8) + s[2, 1]}};
```

```
detjuA = FullSimplify[Det[JuA]];
```
vereinfache voll··· Determinante

Here we plug in the solution ueq:

```
detjuAeq = FullSimplify[detjuA /. ueq];
```
vereinfache vollständig

```
detjuAeq=-(r-s[1,1]) (r-s[1,2]) s[1,2] (r-s[1,3]) (s[1,1]-[illegible]
          /(r s[1,1]-2 r s[1,2]+s[1,1] s[1,2]+(r-2 s[1,1]+s[1,2]) s[1,3]) (r s[1,1]+r[illegible]
```

$$-\left(\left((r-s[1,1])\,(r-s[1,2])\,s[1,2]\,(r-s[1,3])\,(s[1,1]-s[1,3])^2\,(r-s[2,1])\,s[2,1]\right)\Big/\left((r\,s[1,1]-2\,r\,s[1,2]+s[1,1]\,s[1,2]+(r-2\,s[1,1]+s[1,2])\,s[1,3])\,(r\,s[1,1]+r\,s[1,3]-2\,s[1,1]\,s[1,3]+(-2\,r+s[1,1]+s[1,3])\,s[2,1])\right)\right)$$

```
FullSimplify[
vereinfache vollständig
 detjuAeq == -(r^2 (r - s[1, 2]) (s[1, 1] - s[1, 3])^2 (r - s[2, 1]))/((r - s[1, 1]) (r - s[1, 3])) p[1] q[1] /. ptr /. qtr]
True
```

$$\text{detjuAeq} == -\frac{r^2 (r - s[1,2]) (s[1,1] - s[1,3])^2 (r - s[2,1])}{(r - s[1,1]) (r - s[1,3])} p[1]\, q[1] \;/.\; ptr \;/.\; qtr$$

The forefactor is always negative because r-s[i,j]>0 ∀ i,j and r>0. p[1] and q[1] are positive. Thus the determinant is negative.
